# Loss of Cdc13 causes genome instability by a deficiency in replication-dependent telomere capping

**DOI:** 10.1101/757864

**Authors:** Rachel E Langston, Dominic Palazzola, Erin Bonnell, Raymund J. Wellinger, Ted Weinert

## Abstract

In budding yeast, Cdc13, Stn1, and Ten1 form a telomere binding heterotrimer dubbed CST. Here we investigate the role of Cdc13/CST in maintaining genome stability, using a Chr VII disome system that can generate recombinants, loss, and enigmatic unstable chromosomes. In cells expressing a temperature sensitive *CDC13* allele, *cdc13^F684S^*, unstable chromosomes frequently arise due to problems in or near a telomere. Hence, when Cdc13 is defective, passage through S phase causes Exo1-dependent ssDNA and unstable chromosomes, which then are the source for whole chromosome instability events (e.g. recombinants, chromosome truncations, dicentrics, and/or loss). Specifically, genome instability arises from a defect in Cdc13’s replication-dependent telomere capping function, not Cdc13s putative post-replication telomere capping function. Furthermore, the unstable chromosomes form without involvement of homologous recombination nor non-homologous end joining. Our data suggest that a Cdc13/CST defect in semi-conservative replication near the telomere leads to ssDNA and unstable chromosomes, which then are lost or subject to complex rearrangements. This system defines a links between replication-dependent chromosome capping and genome stability in the form of unstable chromosomes.

## Introduction

Maintaining chromosomes through generations of cellular divisions is an essential process. Telomeres, the highly conserved structures capping the ends of linear chromosomes, act as one safeguard against chromosome, and hence, genome instability. Telomeres consist of repeated G-rich sequence elements ending with a 3’-overhang (Wellinger & Zakian, 2012). One important function of telomeres is to counteract the gradual shortening of the telomeric DNA caused by conventional DNA replication (thereby solving the “end-replication problem” (Gilson & Géli, 2007; Soudet *et al*, 2014)). This is achieved by the telomeric TG repeats becoming a substrate for the conserved reverse transcriptase telomerase, which recognizes and extends the repeat in a highly regulated manner (Wu *et al*, 2017; Greider & Blackburn, 1985). Another primary function of telomeres is to protect chromosome ends from improper recognition and processing as a DNA double-strand break (Wellinger & Zakian, 2012; Jain & Cooper, 2010). Intriguingly, some of the features of the telomeres that act to protect the chromosome end also can act as detriments to chromosome stability. For example, secondary structures formed by the G-rich repeats (Sundquist & Klug, 1989; Paeschke *et al*, 2011; Lopes *et al*, 2011) and many DNA-bound proteins can render DNA replication through the region difficult (Ivessa *et al*, 2002; Goto *et al*, 2015). Long non-coding RNAs bound to the telomeric repeats, dubbed TERRA, can also act as a replication block (Costantino & Koshland, 2018; Rippe & Luke, 2015) and foster recombinatorial repair (Graf *et al*, 2017). In contrast to other difficult-to-replicate regions in the chromosome, disrupted fork progression is particularly challenging in the telomere since there is no option of rescue by an oncoming replication fork. General errors in DNA replication, potentially in difficult to replicate regions, have been linked to chromosome instability, one of the hallmarks of cancer (Macheret & Halazonetis, 2015; Mazouzi *et al*, 2014; Lambert & Carr, 2005; Sogo *et al*, 2002; Groth *et al*, 2010; Burrell *et al*, 2013).

Classically, telomeric DNA binding proteins are thought to be important for maintaining the stability of the telomere itself. In *Saccharomyces cerevisiae*, one such protein is the essential single-stranded DNA binding protein Cdc13. Cdc13 has a high specificity for the terminal telomeric G-strand and can bind the G-rich ssDNA either alone or in a complex with Stn1 and Ten1 (the Cdc13/Stn1/Ten1 heterotrimer CST; (Grandin *et al*, 1997; 2001a; Pennock *et al*, 2001)). CST has functional and structural similarity with the heterotrimeric replication protein A (RPA) and thus has also been dubbed t-RPA (Gao *et al*, 2007). Cdc13/CST facilitates telomere functions via several mechanisms: by facilitating telomerase-mediated telomere elongation (Nugent *et al*, 1996; Evans & Lundblad, 1999); by assisting in telomere processing after replication (Soudet *et al*, 2014); and by “capping” the telomere (Garvik *et al*, 1995; Wellinger & Zakian, 2012; Mersaoui & Wellinger, 2019). “Capping” is a term that likely encompasses at least two functions: a putative post-replication capping function wherein CST binds to the 3’ ssDNA TG overhand and blocks degradation, and a DNA replication-dependent function wherein Cdc13 facilitates semi-conservative DNA replication (even independent of telomerase extension; (Lue, 2018)). CST complexes with similar functions have been identified in fission yeast, plants, and mammals (Martín *et al*, 2007; Surovtseva *et al*, 2009; Miyake *et al*, 2009; Chen & Lingner, 2013). These CST complexes additionally have been implicated in rescuing stalled replication forks in difficult-to-replicate sequences (Stewart *et al*, 2012; Kasbek *et al*, 2013; Chastain *et al*, 2016; Wellinger, 2009). Mutations in hCTC1 (the functional homologue of scCdc13) and hStn1 have been linked to Coats Plus Syndrome in humans (Anderson *et al*, 2012; Simon *et al*, 2016). While many classical telomeropathies are linked to telomerase deficiencies and short telomeres (Opresko & Shay, 2017), the telomere dysfunction found in patients with mutated hCTC1 are consistent with errors in telomeric DNA replication (Chen *et al*, 2013).

In this study, we use Cdc13 and a previously developed chromosome instability assay to explore the link between uncapping telomeres and unstable chromosomes. Using a conditional cdc13 mutant (*cdc13^F684S^*) we show that defects in Cdc13 indeed lead to a high rate of instability, most prominently forming unstable intermediates that eventually are resolved to become recombinants, are lost completely, or remain and are propagated in an unstable form. Taking further advantage of the conditional *cdc13* allele, we show that a cell that completes one S phase at the restrictive temperature forms unstable chromosomes in G2. We also provide evidence that the critical function of Cdc13 needed to keep chromosomes stable is in its DNA replication-dependent function, and not in its putative post-replicative chromosome capping function. Molecular and genetic evidence suggests a key role for ssDNA in forming unstable chromosomes, and that unstable chromosome formation does not require NHEJ nor HR to form, suggesting the unstable chromosome might be linear with a ssDNA end-gap. A possible role for hairpins is suggested because *sae2Δ* suppresses instability. Finally, we show that initially, telomere-linked unstable chromosomes progress to form other, more centromere-proximal rearrangements. The results suggest a model: a Cdc13-defect disrupts replisome function, allowing ssDNA degradation and thus end-gaps on a lagging strand template, which then forms an initial unstable chromosome that progresses to other structures during subsequent cell cycles. We also propose a specific, as of yet untested model involving BIR to repair the ssDNA gaps.

## Results

### Chromosome VII Disome System and Cdc13-associated instability

Figure 1 shows the previously-developed disomic chromosome system in *Saccharomyces cerevisiae* used here to analyze chromosome instability in *cdc13* mutants. This system consists of a haploid cell with an extra-numerary, and therefore nonessential, chromosome VII (Fig 1A; based on Meeks-Wagner and Hartwell 1986, (Meeks-Wagner & Hartwell, 1986; Admire *et al*, 2006)). We detect rearrangements of the extra chromosome by selecting for loss of the *CAN1* gene, located about 25 kb from the left telomere (Fig 1A). *CAN1* cells are sensitive to the drug canavanine (Can^S^); when plated on canavanine plates, cells with an unchanged chromosome retain *CAN1* and die. Alternatively, cells that lose *CAN1* through a chromosome rearrangement are resistant to canavanine (Can^R^) and live. The disome is genetically marked to assist in determining the structure of chromosome changes.

**Fig 1.**
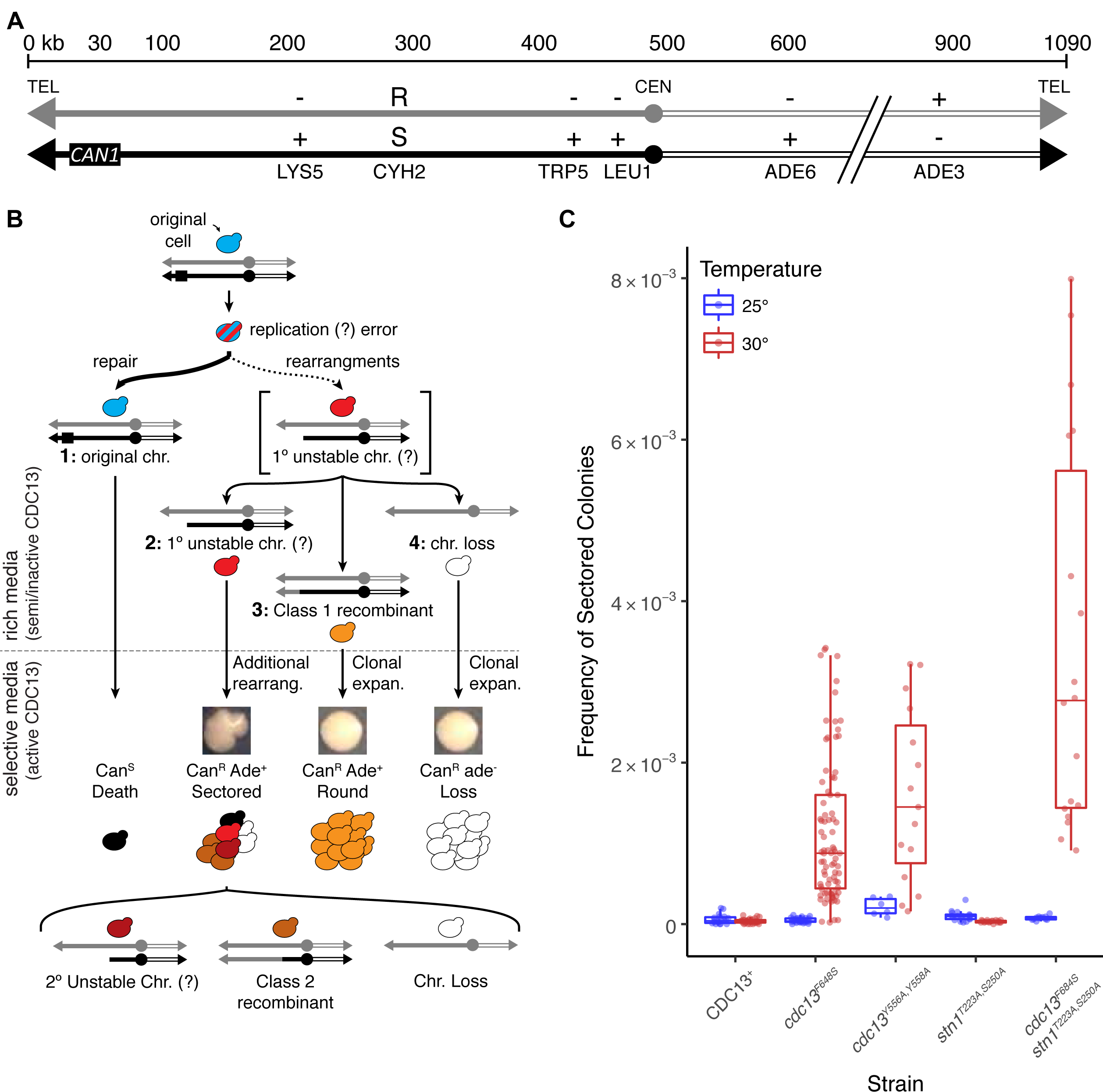
Cdc13 suppresses unstable chromosomes through its activity with CST/t-RPA. **(A)** Chr VII disome system. The two homologues of Chr VII are in black or grey. Position of *CAN1* and 6 heterozygotes are shown. **(B)** Schematic of how diversity arises from one unstable chromosome. An unstable chromosome replicates, then various replicas change again in subsequent cell divisions. Further explanation in text. **(C)** Frequency of unstable chromosomes in *cdc13* mutants from cells grown at 25 °C (blue) or 30 °C (red). The median frequency, IQR, and statistically significant fold changes relative to the Wild Type frequency of the corresponding temperature are noted (* < 0.05; Mann-Whitney U).

In this study, we present evidence that examines a sequence of events shown in Fig 1B (Admire *et al*, 2006; Paek *et al*, 2009). From earlier studies, we proposed that instability might begin at inverted repeats located at a site 403 kb from the telomere. These inverted repeats can fuse and form a dicentric. We subsequently found that deletion of this 403 kb site (a 4 kb deletion) does not affect the frequency of instability: therefore events must begin more telomere proximally (Beyer & Weinert, 2016; Vinton & Weinert, 2017). Furthermore a telomerase mutant induces instability that results in fusion of the inverted repeats in the 403 kb site (Beyer & Weinert, 2016). Knowing events can being in the telomere, and do so in this study, leads us to an overall model shown in Fig 1B. In this model, an initial cell incurs a chromosomal replication error that subsequently gives rise to four different types of cells. Cells can either retain the original chromosome (marked “1”), form an unstable chromosome (“2”), form a recombinant (“3”), or lose the chromosome (“4”). Each outcome has a specific phenotype: unchanged cells die on canavanine, unstable chromosomes form sectored Can^R^ Ade^+^ colonies, recombinants form round Can^R^ Ade^+^ colonies, and loss forms round Can^R^ Ade^-^ colonies. The presence of unstable chromosomes is deduced from the genetic heterogeneity of cells in sectored colonies: cells in the sectored colony have multiple phenotypes, therefore the colony’s founding cell must have had chromosomal instability (Fig 1B). Two additional features of this model are that unstable chromosomes give rise to recombinants and loss, and that there are two classes of recombinants: Class 1 and Class 2. We theorize that Class 1 recombinants arise from the primary unstable chromosome, while Class 2 recombinants develop from either a primary or a secondary unstable chromosome (eg the dicentric at the 403 kb site). The central features of this study are the nature of, and the relationship between, telomeres and unstable chromosomes.

### Chromosomes in cdc13 mutants are unstable at semi-permissive temperatures

To determine the effect of a Cdc13 mutation on chromosome stability, we use a temperature-sensitive conditional mutant, *cdc13^F684S^*(Paschini *et al*, 2012), as Cdc13 is an essential gene. It has been reported that a *cdc13-1* mutation generates recombinants and chromosome loss in both Chr V and Chr VII diploids (unstable chromosomes were not reported; (Carson & Hartwell, 1985; Hartwell & Smith, 1985)). We used the *cdc13^F684S^* allele instead of *cdc13-1* because the former is fully functional at the permissive temperature (Paschini *et al*, 2012), whereas the commonly-used *cdc13-1* alleles is compromised in our cells even at 23 °C (data not shown).

We thus introduced the temperature-sensitive *cdc13^F648S^*allele into the Chr VII disome, grew cells at a permissive (25 °C) or semi-permissive temperature (30 °C) for ∼20 generations, and plated for instability (Fig S1). The *cdc13^F684S^* cells generated an increased frequency of all three types of rearrangements compared to the controls (a *CDC13*^+^ strain grown at 30 °C or *cdc13^F684S^* grown at 25 °C); namely, in the frequency of sectored colonies (Fig 1C), and in recombinants and chromosome loss (Table 1). We tested instability in one other *cdc13* temperature-sensitive mutant (*cdc13^Y556A,Y558A^*) and found it exhibited increased instability as well (Fig 1C; Table 1). We confirmed that the sectored colonies formed in *cdc13*-defective cells contain cells of multiple phenotypes (Fig S2), indicating that, when Cdc13 is defective, unstable chromosomes form and are detected by the sectored colony phenotype. While this study was in progress, we reported elsewhere that a *cdc13^F684S^*mutation also causes unstable chromosomes in a Chr V disome system (Vinton & Weinert, 2017). Thus, cells limited for Cdc13 function generate unstable chromosomes to many if not all chromosomes.

**Table 1.**
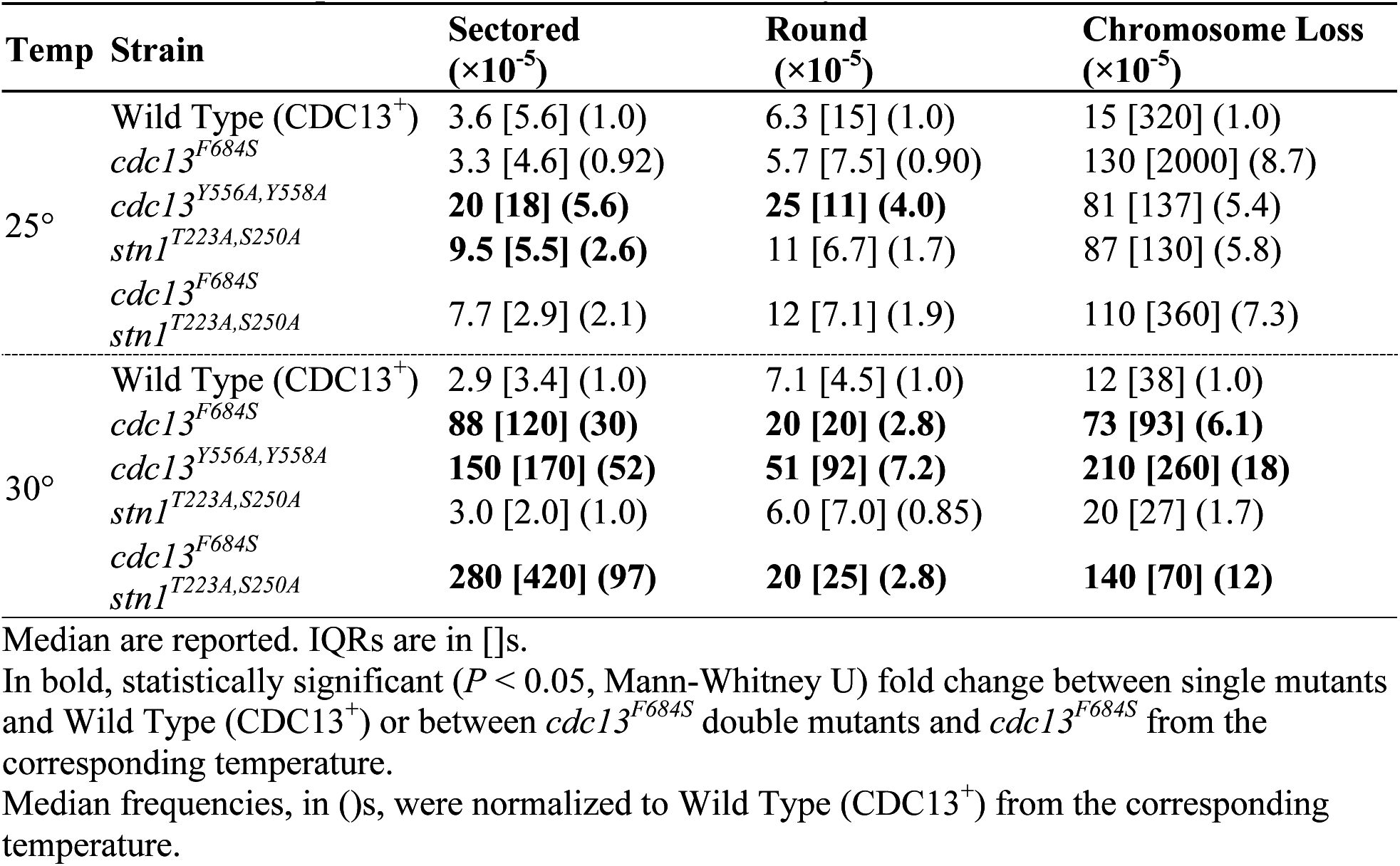
Median frequencies of chromosome instability in *cdc13^F684S^* at 25 and 30 °C

We next determined if Cdc13 prevents chromosome instability alone, or as part of the CST heterotrimeric complex (with Stn1 and Ten1; (Gao *et al*, 2007)). We used a mutation of *STN1*, *stn1^T223A,S250A^*, that partially disrupts Stn1’s ability to concurrently bind Cdc13 and Ten1 (Liu *et al*, 2014). We found that the *stn1^T223A,S250A^* single hypomorphic mutant showed no increase in instability (Fig 1C; Table 1). However, the double mutant *cdc13^F684S^ stn1^T223A,S250A^* generated more sectored colonies than the *cdc13^F684S^* single mutant alone (3.2-fold increase in sectored colonies compared to the *cdc13^F684S^* single mutant, Fig 1C). The frequency of recombinants and loss were not significantly changed from the *cdc13^F684S^* single mutant (Table 1). The simplest explanation for the synergistic increase in sectored colonies in the double mutant is that the CST complex prevents chromosome instability in general, and unstable chromosomes in particular.

### Cdc13 suppresses chromosome instability at the terminal telomere repeat and not at an internal site

Next, we investigated where Cdc13 acts on the chromosome to suppress instability. We expected Cdc13 to act at least at the telomere, yet it could potentially act at any single-stranded sequence (especially a TG-rich sequence) elsewhere on the chromosome (Lin & Zakian, 1996). We took a genetic approach, making use of the fact that the frequency of Can^R^ Ade^+^ round colonies (Class 1 recombinants) is increased by a defect in *cdc13^F684S^* (Table 1), and that Class 1 recombinants likely arise linked to the site of an error (Hackett *et al*, 2001; Hackett & Greider, 2003; Beyer & Weinert, 2016). Thus, if a telomere defect causes recombinants, we expect those recombinants to map to telomere-proximal genetic intervals (as shown previously using *cdc13-1*; (Garvik *et al*, 1995; Carson & Hartwell, 1985)). Indeed, in our Chr VII disome system we found that, in *cdc13*-defective cells, Class 1 recombinants from round colonies were heavily biased towards the telomere-proximal region, compared to events that would be expected given the size of each genetic interval (“random”, Fig 2A). This suggests that a *cdc13*-defect causes errors in or near the telomere (as shown by Garvik et al 1995 for a Chr V strain). Interestingly, the events in *CDC13*^+^ cells are also increased in the telomere-proximal region, suggesting that, while instability is less frequent in *CDC13^+^*cells overall, instability may also originate from the telomere in wild type cells.

**Fig 2.**
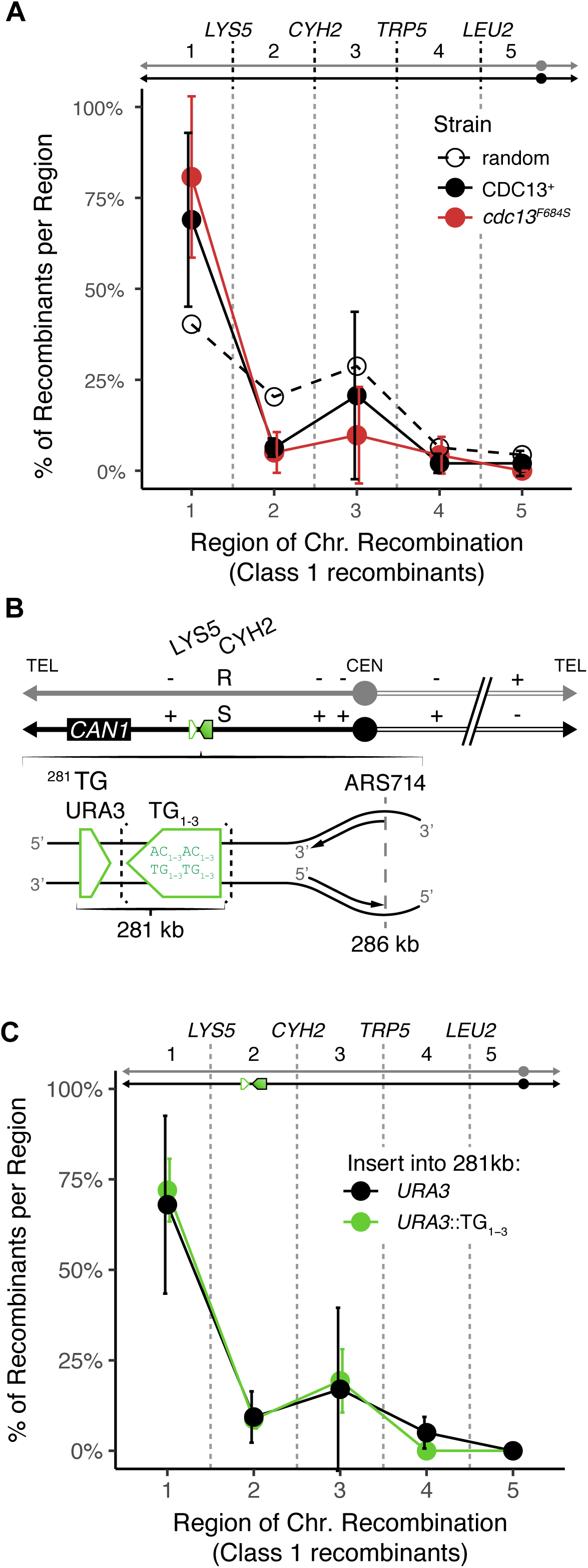
Unstable chromosomes in cdc13^F684S^ form at the telomere, not at internal TG repeats. **(A)** Distribution of Class 1 recombinants from *cdc13^F684S^* round colonies (solid) and the expected distribution (dashed). The average percentage and standard deviation for 4 independent experiments are shown (P = 0.0349; one sample *t* test). **(B)** Schematic for insertion of the TG_1-3_ repeat (green box) into the left arm of Chr VII. The TG_1-3_ insert (green box; bracketed) was marked with *URA3* (green triangle), and was inserted 281 kb from the telomere end. As a control, *URA3* alone was inserted into the same region. **(C)** Distribution of Class 1 recombinants from round colonies from *cdc13^F684S^*with *URA3* or *URA3*::TG_1-3_ inserted into the 281 kb locus. The average percentage and standard deviation for 3 independent experiments are shown.

To address if Cdc13 could suppress instability internally if there were an interstitial telomere-like sequence available, we inserted a 126 bp TG_1-3_ DNA sequence at two different sites distant from the telomere. One site was 281 kb from the telomere (Fig 2B), and another at a locus 15 kb from the centromere (484 kb from the left telomere; Fig S3). Both sequences were inserted such that the TG repeat was oriented relative to nearby origins so that the G-rich strand was on the lagging strand template, mimicking the G-rich strand on the lagging strand template in a natural telomere (Fig 2B). As a control the *URA3* gene alone was inserted into the same regions of the *cdc13^F684S^*strains. These strains were then analyzed for instability, focusing first on the ^281^TG site in *cdc13^F684S^*mutants. We found that the internal TG repeat did not significantly change the frequencies of chromosome instabilities; if instability initiates at this site, the effect is subtle compared to instability elsewhere on the chromosome (Fig S4). To identify any subtle effects by the internal TG repeat, we analyzed the regions of recombination in round colonies (Can^R^ Ade^+^), and found that Class 1 rearrangements from round colonies were still biased to the telomere-proximal region (in both experimental *URA3:TG* or in the *URA3* control strains; Fig 2C); indicating that the internal TG sequence at the 281kb site does not detectably induce events. We made similar observations for the TG repeat inserted 484 kb from the left telomere; there were no changes in the frequency of instability nor in the distribution of recombinants (Fig S3). We therefore conclude that chromosome instability in *cdc13^F684S^* mutants in our system originates only from the telomere end, and not from internal TG repeats of 126 bp. It is important to note that, while the 126 bp interstitial telomeric repeat does not initiate instability, it still influences the progression of instability.

### Cells traversing one cell cycle with inactive Cdc13 are highly unstable

To examine molecular events that underlie instability, we asked if inactivation of Cdc13 in a single cell cycle might generate loss of *CAN1* to form Can^R^ cells. The data reported thus far, in Fig 1C and Table 1, were collected by growing *cdc13* mutant cells at a semi-permissive temperature (where cells delay and resume cell divisions repeatedly for about 20 generations) before being subjected to selection. Thus, we do not know when during these 20 generations events occur, and certainly cannot specify a particular cell cycle phase.

We therefore asked if we could detect instability arising in one cell cycle limited for Cdc13 function. We synchronized *cdc13^F684S^* cells grown at the permissive temperature (25 °C) in early S phase with hydroxyurea (HU); HU-arrested cells have mostly unreplicated and stable chromosomes (Fig 3). We then shifted the HU-synchronized cells to the restrictive temperature (25 °C to 37 °C), washed out the hydroxyurea, and allowed cells to traverse one cell cycle at the restrictive temperature (referred to as the “+HU 25 °C, -HU 37 °C” experiment; Fig 3A). Western blotting verified that the Cdc13^F684S^ protein is quickly degraded when cells are incubated at 37 °C (Fig 3B). Cells completed one S phase and arrested at the *RAD9* G2/M checkpoint (Fig 3C; (Weinert & Hartwell, 1993)). We then reactivated Cdc13^F684S^ by shifting to the permissive temperature (25 °C), allowing cells to repair their chromosomes and resume cell division. We plated cells on selective media to determine if any of these cells had generated altered chromosomes.

**Fig 3.**
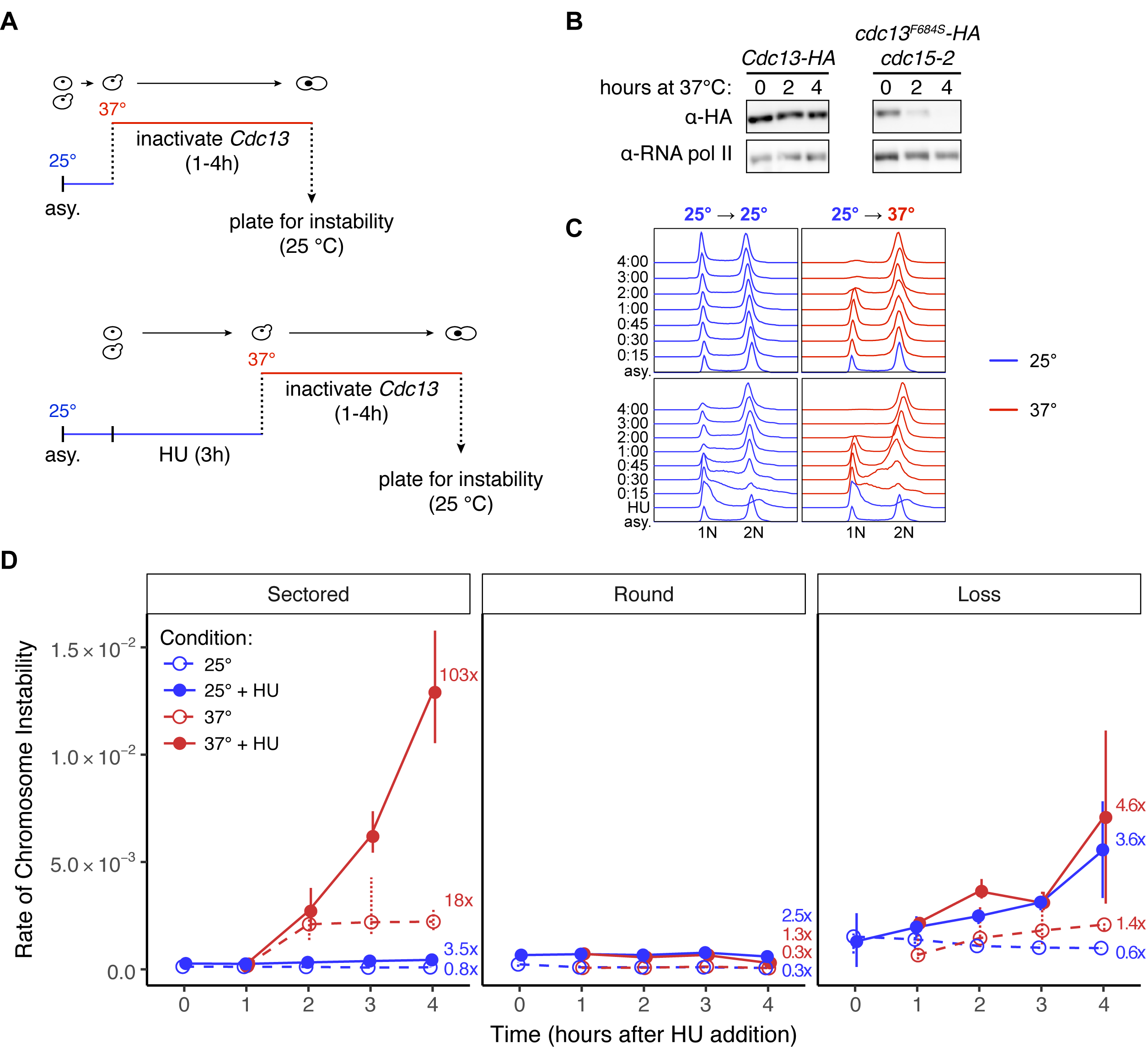
Unstable chromosomes in cdc13^F684S^ form in a single cell cycle. **(A)** The experimental protocol for generating unstable chromosomes in a cell cycle. **(B)** Exponentially growing Cdc13-HA or *cdc13^F684S^-HA cdc15-2* cells were shifted from 25 °C to 37 °C for 2 or 4h. Whole cell extracts were prepared and analyzed by western blot with an anti-HA antibody. Anti-RNA polII was used for a loading control. **(C)** FACS analysis of DNA content. Cells were grown in liquid YEPD and prepared as in 2A. After HU arrest 70% of cells were in G1/S phase. The time increments refer to time after HU wash and shift to either 37 °C or 25 °C. **(D)** The frequency of sectored, round, and loss colonies in *cdc13^F684S^*from 0 to 4 hours after the temperature shift. Blue: incubated at 25 °C; red: incubated at 37 °C. Fold change and significance are calculated from the 0 hour frequency. Data shown are the medians and IQR of 4 independent experiments.

The results from this HU and temperature shift experiment are surprising, and are shown in Fig 3D. Remarkably, cells subjected to HU and then shifted up to 37 °C generated a very high frequency of unstable chromosomes (103-fold increase after 4 hours at 37 °C; Fig 3D). Cells subjected to HU-arrest alone were stable: there was little to no increase in all three types of chromosome changes (+HU, 25 °C; Fig 3D blue). Second, and also surprising, we found that while the +HU 25 °C, -HU 37 °C incubation did increase unstable chromosomes, this regime did not appreciably increase the frequency for either round (Class 1 recombinants) or chromosome loss. Third, shifting to 37 °C without presynchrony in HU increased sectored colonies by 18-fold whereas with presynchrony in HU events were increased 108-fold (and again, only sectored colonies were increased, not round or loss); therefore there appears to be synergy between Cdc13 inactivation and HU treatment.

All together, these results suggest the following. Upon inactivation of Cdc13 and progression through one cycle, cells arrest in G2 with damaged chromosomes. The damaged chromosomes usually repair faithfully (>99% of viable cells are still Can^S^), yet about 1 in 100 cells develop an unstable Chr VII. A surprise is that only sectored colonies are formed (representing unstable chromosomes), but not Class 1 recombinants nor loss; therefore Class 1 recombinants and loss are not generated directly from a Cdc13 defect, but rather arise only from unstable chromosomes (Fig 3D). The +HU 25 °C, -HU 37 °C regime also indicates some interaction with HU-induced DNA replication stall and Cdc13 replication-dependent capping. This experimental paradigm (Fig 3A) provides a rich source of data on Cdc13 function, unstable chromosomes, recombinants, and loss. And, this paradigm allows us to determine when in the cell cycle Cdc13 must be active to preserve chromosome stability.

### Cdc13 must act in S phase, and not in the G2/M phase, to prevent instability

Cdc13 may prevent chromosome instability by any of four mechanisms: facilitating telomerase recruitment, facilitating lagging strand synthesis after telomerase extension, capping the chromosome end (post-replication capping), and/or by facilitating semiconservative DNA replication independent of telomerase function (replication-dependent capping). It is unlikely that Cdc13 must facilitate telomerase activity during one cell cycle to keep chromosomes stable, as a telomerase defect takes many cycles to manifest instability (Hackett & Greider, 2003; Beyer & Weinert, 2016). We therefore suggest that Cdc13’s role in chromosome stability comes from its role in lagging strand synthesis independent of telomerase function (replication-dependent capping) or in chromosome capping (post-replication capping).

To distinguish between these two roles, we made use of the single cell cycle assay described above. We performed two experiments (Fig 4A): in the first, Cdc13 is inactive in both S and in G2/M phases; in the second Cdc13 is active in S phase and inactive in G2/M phase. (A method to inactivate Cdc13 only in S phase has not been developed.) To inactivate Cdc13 in both S and G2/M phases, we repeated the experiment reported in Fig 3. Asynchronously growing cells were shifted from 25 °C to 37 °C for four hours, during which they complete one cell division and arrest in G2/M; Cdc13 is therefore inactive in both S and G2/M phases. The G2/M-arrested cells were then returned to the permissive temperature and plated for instability. As expected from Fig 3D, we detected an increase in unstable chromosomes (and not an increase in loss or Class 1 recombinants; Fig 4D).

**Fig 4.**
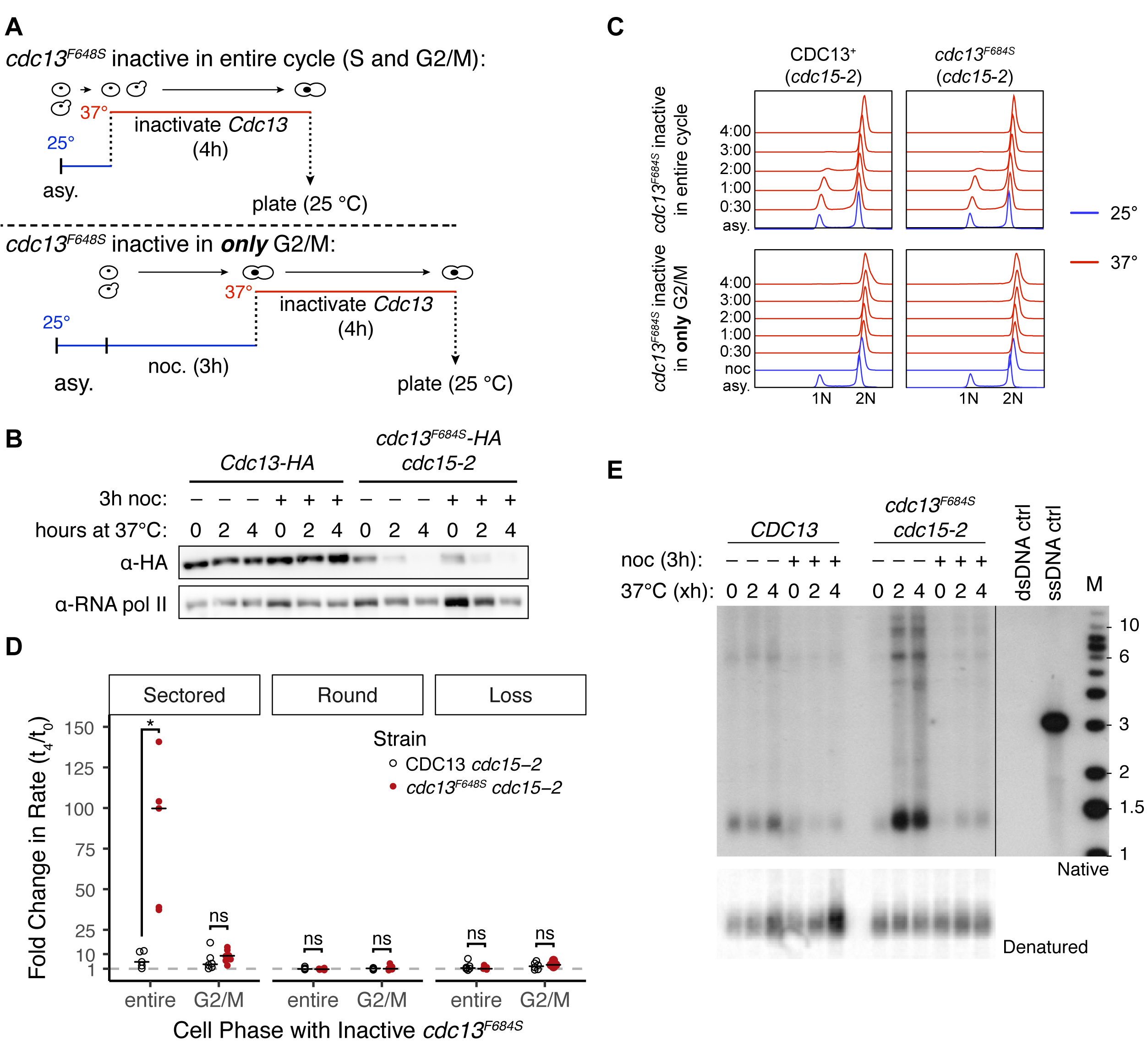
Cdc13 suppresses the formation of unstable chromosomes during S phase. **(A)** The experimental system to either inactivate or activate *cdc13^F684S^*during S phase. *Top:* inactive *cdc13^F684S^* for the entire cell cycle, cells were grown to mid-log then incubated at 37 °C for 4 hours. Aliquots for the instability assay were taken at 0h (before the temp. shift) and after 4 hour of incubation at 37 °C (t_4h_). Bottom: inactivate *cdc13^F684S^*in only G2/M, cells were grown to mid-log at 25 °C and then arrested with nocodazole (noc). After noc was washed out, cells were shifted to 37 °C and kept arrested in G2/M with additional noc. Aliquots for the instability assay were taken at 0h (before the temp. shift) and after 4 hour of incubation at 37 °C (t_4h_). **(B)** Exponentially growing Cdc13-HA or *cdc13^F684S^-HA cdc15-2* cells were shifted from 25 °C to 37 °C for 2 or 4h. Whole cell extracts were prepared and analyzed by western blot with an anti-HA antibody. Anti-RNA polII was used for a loading control. The – noc samples were shown in 3B, and are shown here for comparison. **(C)** FACS analysis of DNA content. Cells were grown in liquid YEPD as depicted in Figure 4A, and cells were taken at the indicated time points for FACS analysis. Blue traces: 25 °C, red traces: 37 °C. After noc arrest 99% of cells were in G2/M. The time increments refer to time after noc removed and the temperature was shifted. **(D)** The average fold change in frequency of unstable chromosomes between t_4_ and t_0_. The relevant genotypes are as follows: open circles: CDC13^+^ control (*cdc15-2*); closed circles: *cdc13^F684S^*(*cdc13^F684S^ cdc15-2*). The fold changes were calculated by dividing each t_4_ frequency by the corresponding t_0_ frequency for that particular sample (n = 5). The average was then calculated (horizontal bar). The statistical significance between strains per experiment is shown (* < 0.05; Mann-Whitney U). **(E)** Non-denaturing in-gel hybridization using a CA oligonucleotide probe with XhoI-digested DNA from the indicated samples (*top*). To control DNA input, the gel was denatured and transferred to a nitrocellulose membrane and hybridized with a telomeric probe (*bottom*).

The second, more elaborate experiment determines the fate of chromosomes in cells where Cdc13 was active in S phase and inactive in the G2/M phase. We grew *cdc13^F684S^* cells at 25 °C, allowed them to progress normally through S phase, and then arrested them in G2/M at 25 °C with the spindle poison nocodazole. We then shifted the cells to the restrictive temperature (37 °C) for four hours while keeping the cells in nocodazole (thus Cdc13 is inactive in G2/M). We confirmed that the Cdc13^F684S^ protein is equally unstable at 37 °C in both treatments (Fig 4B). We also verified using flow cytometry and nuclear staining (DAPI) that most cells (∼80%) did not escape the nocodazole arrest during the four hours at high temperature, though ∼20% of cells did escape (Fig 4C, S5). We therefore introduced a *cdc15-2* mutation that causes an arrest in telophase to prevent these ∼20% escaping cells from progressing through to the next cell cycle; Fig S5 (Hartwell *et al*, 1974; Jaspersen *et al*, 1998). We then shifted the G2/M arrested cells from the restrictive to the permissive temperature to reactive Cdc13^F684S^, washed out the nocodazole, and plated cells for instability. Remarkably, there was no increase in the frequency of sectored colonies (nor round, nor loss; Fig 4D) in this “inactive in G2/M phase only” experiment. We conclude that inactivation of *cdc13^F684S^* in S phase causes instability, but inactivation in G2/M does not; thus it is Cdc13’s function during DNA replication that is critical for chromosome stability, not its role in post-replication chromosome capping.

We also implemented this *cdc13*-inactivation in a version of the “G2/M-only” strategy using arrest at the *cdc15-2* telophase arrest point. We grew cells at 25 °C, treated them with nocodazole at 25 °C, then shifted the cells to the restrictive temperature after washing out nocodazole, whereupon cells exit metaphase and arrest in telophase. After 4 hours at 37 °C, telophase-arrested cells were shifted to 25 °C and assessed for instability. Again, we found that inactivating *cdc13* in G2/M and in telophase did not cause an increase in instability (“T” arrest; Fig S5).

### Inactivation of cdc13^F684S^ in S phase, but not in G2/M, causes extensive ssDNA in telomeres

A Cdc13-defect is known to generate telomere-proximal ssDNA, wherein the 5’ strand is degraded, leaving the 3’ TG rich strand unpaired. A plethora of studies have examined this so-called “chromosome capping” role of Cdc13 (Nugent *et al*, 1996; Garvik *et al*, 1995; Grandin *et al*, 2001b; Grandin & Charbonneau, 2003; Booth *et al*, 2001; Ngo & Lydall, 2010; Vodenicharov & Wellinger, 2006). However, it remains unclear whether the ssDNA arises due to a defect in DNA replication as cells progress through S phase, or a defect in post-replication chromosome capping. As instability arises only when Cdc13 is defective during S phase, we wished to determine if ssDNA arises only during S phase as well. We therefore measured ssDNA in telomeres when Cdc13^F684S^ was defective in S and G2/M phase, versus defective in G2/M only, using the well-established native in-gel hybridization detection method ((Dionne & Wellinger, 1996; Vodenicharov & Wellinger, 2006); see Methods). We detected extensive ssDNA when Cdc13 was inactive in S and G2/M phase, and very little ssDNA when Cdc13 was inactive in G2/M alone (Fig 4E). Thus, generation of ssDNA occurs primarily as cells transit S phase when Cdc13 is defective, and formation of ssDNA correlates with formation of unstable chromosomes. (We note that four studies have detected ssDNA in G2/M arrested cells, though only the first two cited compared the amount of ssDNA generated during S phase versus in G2/M (Diede & Gottschling, 1999; Ngo & Lydall, 2010; Vodenicharov & Wellinger, 2006; Hirano & Sugimoto, 2007). Comparative studies also used nocodazole to prevent cell cycle progression at the high temperatures. Indeed, we find that *cdc13^F684S^ CDC15*^+^ cells shifted to 37 °C with nocodazole generated ssDNA, most likely due to some cells escaping the nocodazole arrest; Fig S6).

### An unexpected phenotypic lag of Can^S^ probably enables the single cell cycle study

Several features of our Chr VII disome system suggested to us we would not detect loss of *CAN1* gene in one cell cycle, even if it were to occur, yet we do. There are three issues that concern the *CAN1* gene. First, the gene is 25 kb from the telomere; a telomere defect due to Cdc13 dysfunction must inactive the *CAN1* gene many kilobases away from the telomere within several hours. Second, the gene may only be gone from one of the two sister chromosomes; a G2-arrested cell may have one *CAN1* gene still intact, while its sister may have lost its *CAN1* gene. And finally, even in the absence of the *CAN1* gene, a phenotypic lag may persist until the Can1 protein is cleared from the cell surface. How could Can^R^ cells have been generated in one cell cycle? We find that the death of Can^S^ cells has a phenotypic lag. Can^S^ cells on canavanine sustain 1 or 2 cell divisions, though slowed ones (Fig S7), and remain alive for several hours. We surmise that while the uptake of canavanine is eventually crippling to a Can1^+^ cell, cells do not die immediately (Fig S7). We therefore speculate that the lag of cell death may allow cell division, to form *CAN1*^+^ and *can1^-^* daughter cells, and allow clearing of the Can1 protein in the *can1^-^* cell. Whatever the exact mechanism of *CAN1* inactivation, lack of Cdc13 causes chromosome damage in the first cycle that manifests into a Can^R^ cell.

### Genetic analyses also suggest that ssDNA causes instability

We next used genetics to gain further insight into how unstable chromosomes are formed. We measured the frequency of instability in a series of double mutants (*cdc13^F684S^ mutX*) to identify possible synergistic, suppressive, or epistatic interactions. We grew single and double mutants at the semi-permissive temperature (30 °C) for about 20 generations, then determined instability by plating on selective media. We found that three mutations suppressed the formation of unstable chromosomes (sectored colonies), including mutations in the nucleases Exo1 and Sae2, and in the helicase Pif1 (Table 2, Table S1). Suppression by *exo1Δ* was the most robust. Curiously, while an *exo1Δ* mutation led to a decrease in sectored Can^R^ Ade^+^ colonies, it led to a concomitant increase in round Can^R^ Ade^+^ Class 1 recombinants. We infer that ssDNA formed during an S phase defect is important for instability, and when limited by an *exo1* mutation, the initial lesion is converted into a Class 1 recombinant instead of an unstable chromosome.

**Table 2.** Median frequencies of chromosome instability in *cdc13^F684S^* mutants

|  | **Strain** | **Sectored**<br>**($\times 10^{-5}$)** | **Round**<br>**($\times 10^{-5}$)** | **Chromosome Loss**<br>**($\times 10^{-5}$)** |
| --- | --- | --- | --- | --- |
|  | Wild Type<br>(CDC13<sup>+</sup>) | 2.9 [3.4] (1.0) | 7.1 [4.5] (1.0) | 12 [38] (1.0) |
|  | *cdc13<sup>F684S</sup>* | **88 [120] (30)** | **20 [20] (2.8)** | **73 [93] (6.1)** |
| Nucleases | *exo1Δ* | 11 [13] (3.8) | **48 [15] (6.8)** | 44 [100] (3.7) |
|  | *cdc13<sup>F684S</sup> exo1Δ* | **5.7 [3.8] (2.0)** | **380 [69] (54)** | **20 [21] (1.7)** |
|  | *sae2Δ* | 7.3 [2.6] (2.5) | 10 [7.7] (1.4) | 32 [33] (2.7) |
|  | *cdc13<sup>F684S</sup> sae2Δ* | **35 [16] (12)** | 8.9 [5.5] (1.3) | 89 [55] (7.4) |
| RPA mutants | *rfa1-t33* | **260 [96] (90)** | **20 [13] (2.8)** | **1300 [450] (110)** |
|  | *cdc13<sup>F684S</sup> rfa1-t33* | **270 [55] (93)** | **10 [4.6] (1.4)** | **870 [390] (73)** |
|  | *rfa1-t33 sae2Δ* | **130 [17] (45)** | **3.9 [3.8] (0.55)** | **910 [260] (76)** |
| HR | *rad52Δ* | 16 [4.7] (5.5) | < 1 | 14 [11] (1.2) |
|  | *cdc13<sup>F684S</sup> rad52Δ* | **230 [320] (79)** | < 1 | **1200 [1500] (100)** |
| NHEJ | *lig4Δ* | 2.8 [2.1] (1.0) | 8.6 [6.9] (1.2) | 12 [13] (1.0) |
|  | *cdc13<sup>F684S</sup> lig4Δ* | 74 [120] (26) | 16 [30] (2.3) | 12 [31] (1.0) |
| RPA mutants | *rfa1-t33* | **260 [96] (90)** | **20 [13] (2.8)** | **1300 [450] (110)** |
|  | *cdc13<sup>F684S</sup> rfa1-t33* | **270 [55] (93)** | **10 [4.6] (1.4)** | **870 [390] (73)** |
|  | *rfa1-t33 sae2Δ* | **130 [17] (45)** | **3.9 [3.8] (0.55)** | **910 [260] (76)** |
| DNA damage<br>checkpoint | *rad9Δ* | **54 [35] (19)** | 8.0 [4.6] (1.1) | 35 [130] (2.9) |
|  | *cdc13<sup>F684S</sup> rad9Δ* | **280 [330] (97)** | 12 [8.2] (1.7) | **960 [1100] (80)** |

Median values and IQR, in [], are reported.
In bold, statistically significant ( $P < 0.05$ ; Mann-Whitney U) fold change between single mutants and Wild Type (CDC13<sup>+</sup>) or between *cdc13<sup>F684S</sup> mutX* and *cdc13<sup>F684S</sup>* (except for *rfa1-t33 sae2Δ*, which is in relation to *rfa1-t33*).
In ()s, median frequencies were normalized to Wild Type (CDC13<sup>+</sup>).

We tested for a role of a second nuclease, Sae2, known to cleave hairpins and degrade 5’- 3’ from a double strand break (Lengsfeld *et al*, 2007; Lobachev *et al*, 2002; Cannavo & Cejka, 2014). We were particularly interested in Sae2 because Deng et al showed that a *sae2* mutation enhanced instability in a *rpa1-T33* mutant in a Chr V GCR assay (Deng *et al*, 2015). Their interpretation of *sae2* enhancement was that extensive ssDNA of a *rpa1-T33* mutant leads to a foldback hairpin structure which, when not cleaved by Sae2, leads to instability via replication of the uncleaved hairpin and the formation of a dicentric chromosome. We therefore tested if extensive ssDNA in *cdc13* mutants might form unstable chromosomes by a similar *SAE2*- sensitive foldback, hairpin mechanism. However, we found that *cdc13^F684S^ sae2Δ* mutants had a *lower* frequency of instability than *cdc13^F684S^*. We also asked if Sae2 might suppress instability of Chr VII disome in an *rpa1-T33* mutant, as Sae2 suppresses instability in the Deng et al Chr V system. We found that *rpa1-T33* mutants had a high frequency of instability in our Chr VII (as in the Chr V system), but, in contrast to Deng et al, Sae2 enhanced Chr VII instability whereas Sae2 suppressed instability in the Chr V system (in other words *rpa1-T33 sae2Δ* had a *lower* level of instability than *rpa1-T33 SAE2* (Table 2)). Therefore it is unlikely that the hairpin mechanism proposed by Deng et al for their Chr V system is responsible for Chr VII’s instability. However, hairpins could still play a role in our system. For example, Sae2 nicking at a hairpin may assist Chr VII system instability.

### Neither NHEJ nor HR are involved in initiation of instability, and the RAD9 checkpoint suppresses instability

We generated numerous other *cdc13^F684S^ mutX* double mutants to probe mechanisms of instability. We tested the involvement of NHEJ and HR, motivated by the idea that sisters might need to fuse to form the initial unstable chromosomes. We found that neither *cdc13^F684S^ lig4Δ* nor *cdc13^F684S^ rad52Δ* double mutants had altered frequencies of unstable chromosomes relative to the corresponding single mutants (Table 2), therefore neither Lig4 nor Rad52 are needed to form unstable chromosomes (as we had concluded in an earlier study of instability arising spontaneously; see (Admire *et al*, 2006; Paek *et al*, 2009)). We did find that Class 1 recombinants are dramatically reduced in *cdc13^F684S^ rad52Δ* double mutants, as one might expect if HR is involved in recombination between homologs.

We additionally tested many other mutations seeking insight into mechanisms. The only mutants with increased instability greater than 2-fold were *tel1Δ* and *rad9Δ*; the *cdc13* and *tel1Δ* interaction appears to be additive, while the *cdc13* and *rad9Δ* interaction is more synergistic (a 5- fold increase in instability over either single mutant; Table 2, S1). Cdc13 and Tel1 may therefore act independently, while Rad9 may suppress instability by either limiting degradation by suppressing Exo1 (Lydall & Weinert, 1995; Zubko *et al*, 2004) by suppressing cell division of cells with an unstable chromosome, or by both mechanisms. Notably, Cdc13 and Rfa1 appear epistatic, indicating that they are in the same pathway when it comes to protecting the cell from instability.

### Progression of telomere-proximal unstable chromosomes

Finally, we wished to gain further insights into early events by characterizing the fates of the telomere-proximal unstable chromosome. We have shown already that telomere-proximal unstable chromosomes in a *cdc13^F684S^* mutant generate Class 1 recombinants enriched near the telomere, as well as chromosome loss (Fig 2A, Table 1). Additionally, we tested if the telomere-proximal unstable chromosomes also progress to form the previously characterized dicentric at the 403 kb locus (Admire *et al*, 2006; Paek *et al*, 2009). To detect this dicentric we employed a genetic strategy developed previously called the “*URA3* proxy system” (Fig 5A; (Paek *et al*, 2009)). In brief, in a normal Chr VII, an inverted repeat centered 403 kb from the telomere undergoes two consecutive fusions reactions, the first one between inverted repeats to form a dicentric and the second between direct repeats to resolve the dicentric to a monocentric isochromosome. The *URA3* proxy system recapitulates these two reactions, forming a dicentric and then a monocentric isochromosome, but using DNA fragments generated from the *URA3* gene such that completion of the two reactions generates an intact *URA3* gene in what was a *ura3*^-^ cell, and thus generating a Ura^+^ cell (Fig 5A). We generated a *cdc13^F684S^* mutant with the appropriate *URA3* fragments (which did not alter the frequencies of instability; Fig 5B), and analyzed sectored colonies formed at 30 °C; Ura^+^ cells indeed arose about 1,000 fold more frequently in sectored colonies (founded by a telomere-proximal unstable chromosome) than in controls (round colonies founded by a stable Class 1 recombinant; Fig 5C). We conclude that a telomere-proximal unstable chromosome can progress to form a centromere-proximal dicentric, which resolves to an isochromosome. Furthermore, if indeed the unstable chromosome was telomere-proximal, we predicted that each sectored colony would contain some Lys^+^ cells (indicative of Class 2 recombinants near the telomere) as well as Ura^+^ cells. Indeed, we detected Lys^+^ and Ura^+^ cells from each of 10 sectored colonies analyzed. We conclude that telomere-proximal unstable chromosomes in *cdc13* mutants progress to form Lys^+^ recombinants as well as the 403 kb dicentric, and subsequently the isochromosome.

**Fig 5.**
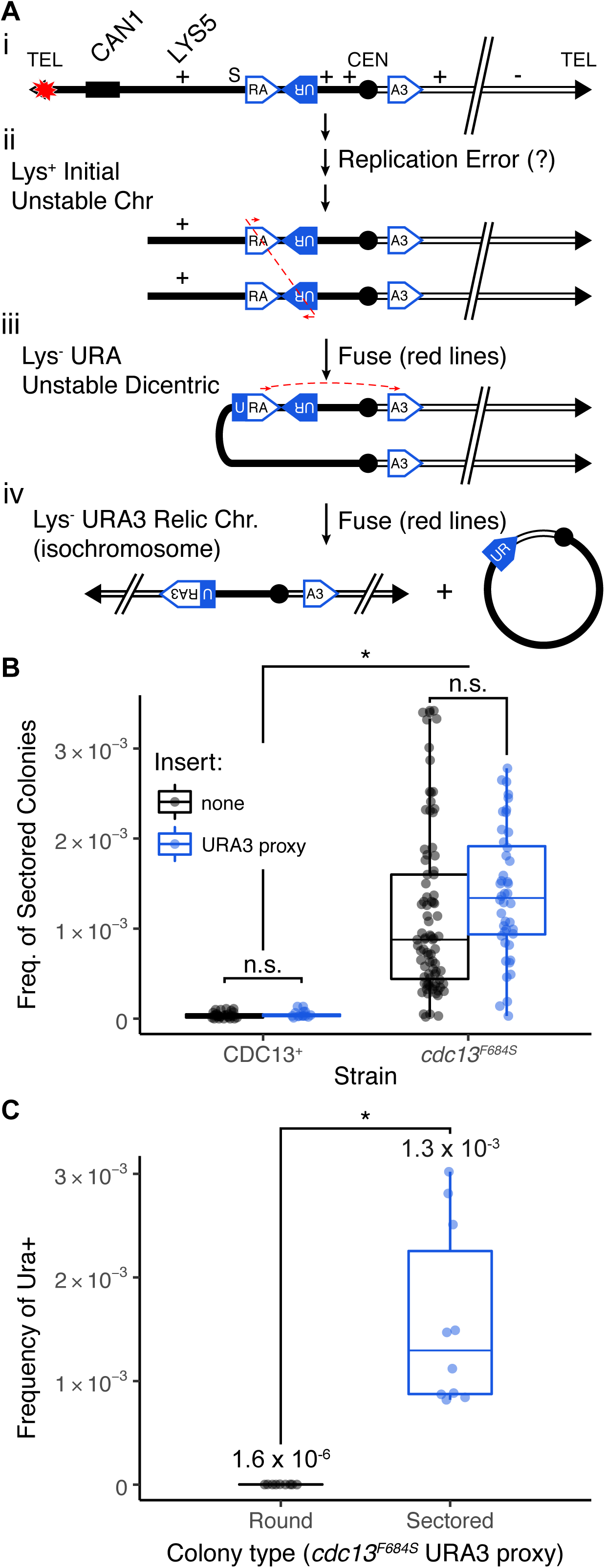
cdc13^F684S^-induced unstable chromosomes progress to centromere-proximal dicentric. **(A)** Model for the URA3 proxy translocation. Two modules, RA and RU are inserted into the left arm as inverted repeats, and the third module (A3) is inserted into the right arm (i). An inverted fusion between U<u>R</u> and <u>R</u>A forms a dicentric chromosome (ii). A secondary event, a direct repeat fusion between UR<u>A</u> and <u>A</u>3 (iii), removes a centromere and creates a functional *URA3* gene (iv). **(B)** Frequency of sectored colonies between strains with (blue) and without (black) the *URA3* proxy system is unchanged. Fold change and statistical significance relative from the corresponding Wild Type frequency are noted (* < 0.05; Mann-Whitney U). **(C)** Average frequency of *URA3*^+^ cells in round (black) and sectored colonies (blue) in *cdc13^F684S^*. The average frequency of Ura^+^ and the standard deviation are shown. The statistical significance between round and sectored colonies is depicted (* < 0.05; Mann- Whitney U).

### Class 1 and Class 2 recombinants

In studying unstable chromosome progression, we discovered a rearrangement present in strains that contain the 126 base pair TG repeat. Specifically, when the 126 base pair TG repeat (see Fig 2B) was inserted into a site 281 kb from the telomere, we found no increase in Class 1 recombinants linked to this site, but did detect a dramatic increase of Class 2 recombinants linked to this site (Fig 6). We analyzed these Class 2 recombinants by pulse field gels and found that most contained a chromosome truncation corresponding in length to truncation at the inserted TG repeat (Fig S8). As expected, we did not detect an increase in Class 2 recombinants when we inserted the TG repeat in the opposite orientation, which could not act as a seed for *de novo* telomere addition (Fig S9). We conclude that a *cdc13* defect in the telomere leads initially to Class 1 recombinants near the telomere, and then later to truncation events hundreds of kilobases from the telomere end (Class 2 recombinant). The insertion of the TG repeats allowed for these truncations to be stabilized.

**Fig 6.**
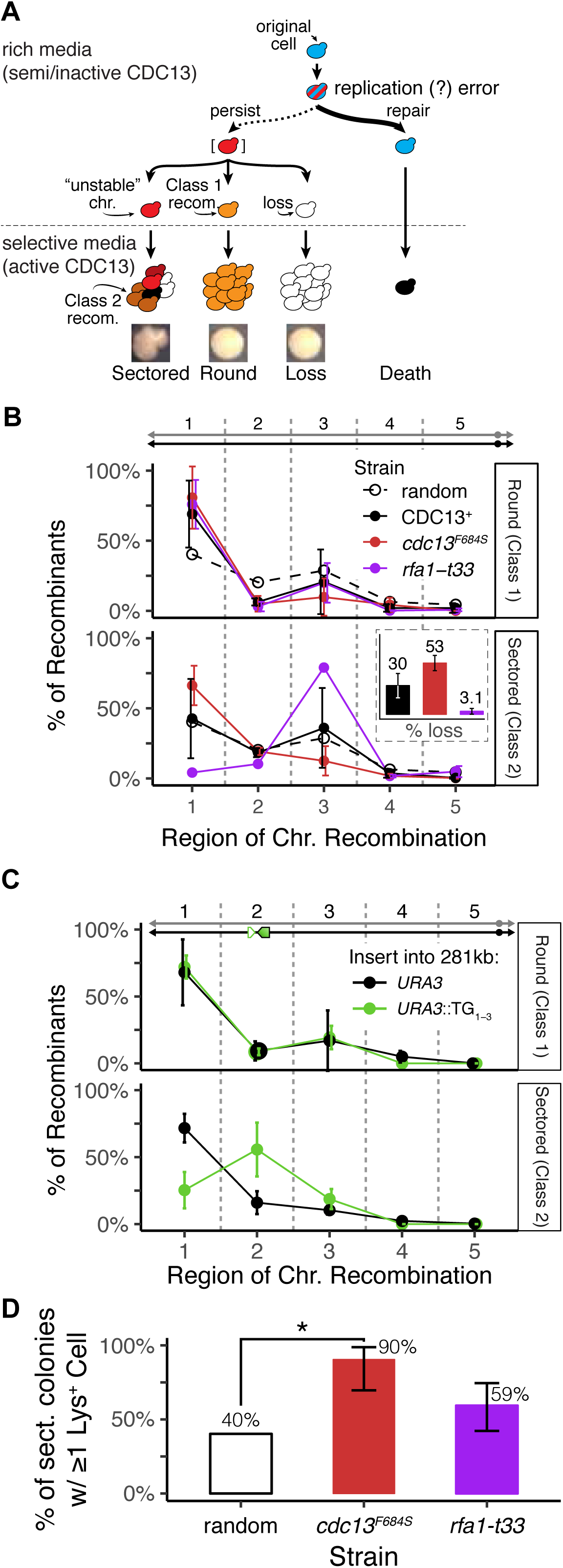
Class 1 and Class 2 recombinants. **(A)** Schematic of how diversity arises from one unstable chromosome. Simplified from 1B. **(B)** Distribution of genotypes from round (Class 1 recombinants) and sectored colonies (Class 2 recombinants) from *cdc13^F684S^* (black) or *rfa1-t33* (purple). The expected distribution (“random”) is dashed. The average percentage and standard deviation for 3 independent experiments are shown. *Inset*: the frequency of chromosome loss with in the sectored colony for *cdc13^F684S^* (black) and *rfa1-t33* (purple). **(C)** Distribution of Class 2 recombinants from sectored colonies from *cdc13^F684S^*with *URA3* or *URA3*::TG_1-3_ inserted into the 281 kb locus. The average percentage and standard deviation for 3 independent experiments are shown. **(D)** Percentage of sectored colonies that retained at least 1 Lys^+^ cell for *cdc13^F684S^* and *rfa1- t33*. The expected distribution (“random”) is white.

The distinction between Class 1 and Class 2 recombinants is not unique to the ^281^TG containing strains. In some strains Class 1 and Class 2 recombinants have similar profiles (Fig 6B; *cdc13^F684S^*), but in other strains the profiles differ dramatically (*rfa1-T33*, Fig 6B; *cdc13^F684S^*^281^TG, Fig 6C). We suggest that in *cdc13* mutants alone, the early unstable chromosomes do not progress before selection, thus Class1 and 2 recombinants have similar profiles. In *cdc13^F684S^* ^281^TG strains, either progression is more rapid, or recombination between the telomere and the ^281^TG site may manifest slowly, generating two different profiles. Finally, *rpa1-T33* mutants also exhibit different profiles; perhaps progression is rapid for unstable chromosomes on selective media. Whatever the mechanisms, the distinction between Class 1 and Class 2 recombinants suggests the unstable chromosomes are very dynamic.

## Discussion

Here we report how chromosome instability is caused by a defect in the telomere-binding protein Cdc13, acting in the CST complex. First, we showed that instability is due to a defect in Cdc13/CST, and instability is linked to the TG repeats at the *bonafide* telomere and not to a TG repeat inserted internally (Fig 1, 2). Events linked to the internal TG repeats do arise, but we find they arise from a previously-formed telomere-linked unstable chromosome as Class 2 recombinants (Fig 6). Second, the unstable chromosomes originate from damaged chromosomes containing ssDNA (Fig 4), and the ssDNA is generated from a mutant Cdc13-associated S phase defect and degradation by Exo1 (Table 1). In addition, instability is not correlated with a defect in the so-called post-replication end-capping function of Cdc13 (Fig 4; Table 1). Third, instability is suppressed by *sae2* in both *cdc13^F684^* and *rpa1-T33* mutants (Table 1), suggesting a role for cleavage of hairpins. Fourth, the initial telomere-linked unstable chromosomes progress to form Class 1 recombinants, simple loss, a chromosome truncation, a known dicentric at the 403 site, and centromeric-unstable chromosomes that resolve to form Class 2 recombinants (Fig 5, 6). We discuss key features of our findings, followed by a speculative model.

### The nature of Cdc13 function and a replication defect

Clearly understanding the intricacies of Cdc13 function is key to understanding instability. Cdc13 acts in a heterotrimer with Ten1 and Stn1. Because a *cdc13^F684S^ stn1^T223,S250A^* double mutant is more unstable than either single mutant, it is likely the heterotrimer that needs to function to suppress instability. What function of Cdc13/CST is relevant here? Interaction with telomerase is unlikely to be involved, as a telomerase defect takes many generations to cause instability (Hackett *et al*, 2001; Beyer & Weinert, 2016), whereas a defect in Cdc13 takes one generation to cause instability (this study).

A DNA replication defect in *cdc13* mutants is likely important, as we find that inactivation of Cdc13 in S phase causes instability, but not inactivation in G2/M cells (Fig 4). That inactivation of Cdc13 in G2/M cells does not generate ssDNA or cause instability calls into question the extent to which Cdc13/CST is really capping the chromosome end independent of replication. We suggest that during replication, CST could assist lagging-strand synthesis by acting at either the end of the chromosome or at the replication fork. We favor the hypothesis that Cdc13 acts from the end of the chromosome rather than the replisome since we do not detect chromosome instability originating from the interstitial 126 bp TG repeat. We do find some interaction between HU treatment and *cdc13* in instability, implying a defect in DNA replication causes instability; but how HU treatment affects CST function at the telomere is unclear. Some telomere proteins may act at the fork to facilitate progression; we note that Taz1 in fission yeast is required to prevent stalled forks in single copy sequence immediately adjacent to the telomere repeats, suggesting Taz1 may facilitate fork movement through the repeats (Miller *et al*, 2006).

### What happens to a damaged chromosome when Cdc13 is reactivated?

The fate of damaged chromosomes when Cdc13 is reactivated is a question central to understanding the generation and outcome of unstable chromosomes. There are few studies of how Cdc13-defective and arrested cells recover (Maringele & Lydall, 2002). We note that mutants in PRR (e.g. *rad18*Δ) do not synergize with a *cdc13* defect, suggesting that whatever ssDNA gap structures are present are not repairable by PRR. Whether an unstable chromosome is formed before Cdc13 reactivation, or after, is also unclear. The model below posits that extensive ssDNA may engage strand-invasion reactions, not permitting completion of failed lagging strand synthesis.

### Before the Model: Issues regarding CAN1 and Exo1-dependent degradation

There are several issues concerning the position of *CAN1* and degradation by Exo1 to generate unstable chromosomes that affect possible models of instability. Defects in Cdc13 cause extensive degradation that is Exo1 dependent; how far does the degradation proceed, and is the *CAN1* gene rendered single stranded? If Exo1 degrades dsDNA to ssDNA at about 4 kb/hour (Booth *et al*, 2001), and given that *CAN1* resides 25 kb from the telomere, it is unclear to what extent the *CAN1* gene becomes single-stranded in 4 hours in that first cell cycle. Second, we posit that in one cell cycle, only one of the two sisters might lose integrity of the *CAN1* gene. It seems reasonable to posit that one sister chromosome (product of leading strand synthesis) is fully intact when Cdc13 is defective; though this is purely speculative. Apparently, there is sufficient phenotypic lag to enable generation of a Can^R^ and a Can^S^ cell from one cell division. And finally: what occurs in *cdc13^F684S^ exo1* mutant cells to possibly convert an unstable chromosome to a recombinant (Table 1)? The more limited ssDNA gaps in *exo1* mutants seem to favor recombination. The position of *CAN1* relative to the telomere, rate of Exo1-dependent degradation, the fate of sister chromosomes, and the role of smaller versus larger gaps are issues remaining to be resolved to permit a clearer model.

### A Model of Cdc13-induced instability and an unstable chromosome

Despite the uncertainties of *cdc13*-induced instability, we know enough to propose the specific model in Figure 7, based on our findings and observations on Cdc13 function. In this model, a defect in CST post replication has no consequences for chromosome stability. A defect in CST during S phase, however, generates one intact and one defective sister chromosome if one strand is intact (leading strand) and one defective (lagging strand (circled); (Lue, 2018; García-Rodríguez *et al*, 2018)). The damaged sister attempts repair by strand invasion and replication. If repair occurs using the intact sister (not shown), no genetic change arises (unless by increased point mutation in the *CAN1* gene). If repair occurs from the homolog, the outcomes depend on what sequences pair during strand invasion. Invasion telomere-proximal to *CAN1* would result in no genetic change (not shown). Invasion centromere-proximal to *CAN1* would result in loss of *CAN1* and a Can^R^ cell. Invasion centromere-proximal to *CAN1* might benefit from ssDNA forming hairpins, cleavable by Sae2, forming an internal “nick”. After invasion, either replication proceeds to the telomere to form a recombinant, or replication aborts as second end capture fails. With no second end capture possible, the unstable chromosome undergoes progressive degradation, repeated attempts of strand-invasion and dissociation, forming other forms of chromosome products.

**Fig 7.**
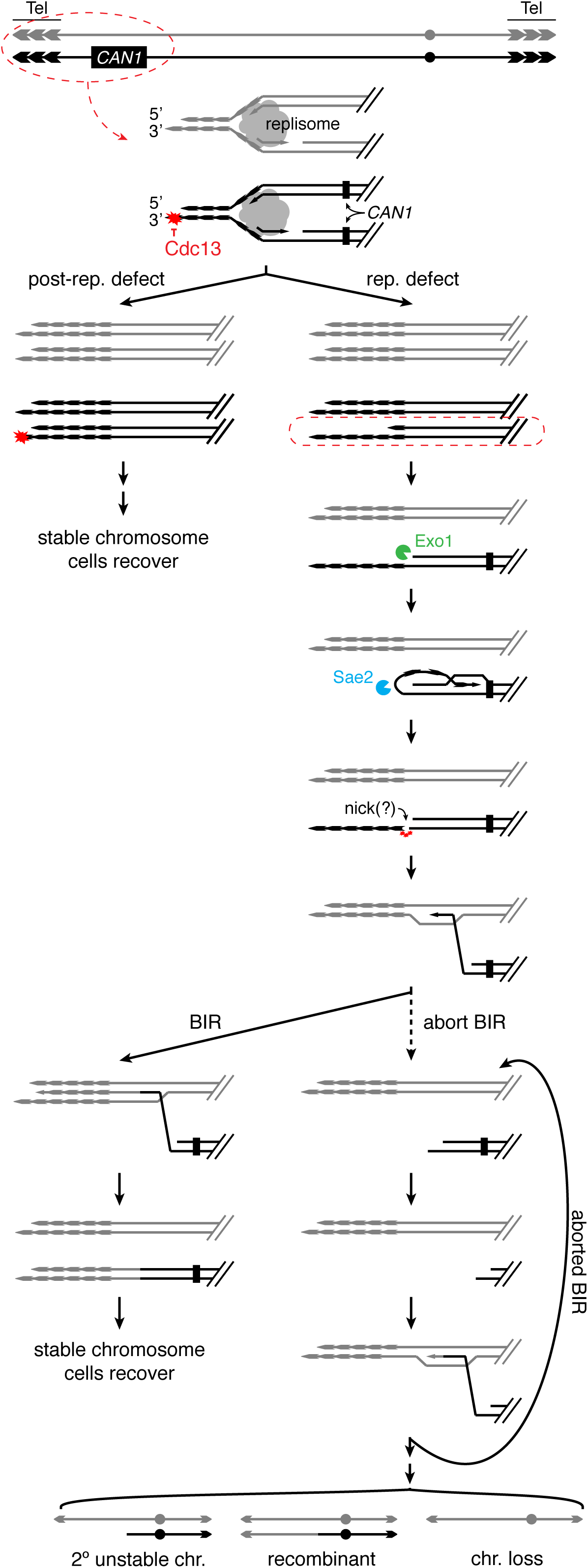
Model for formation of unstable chromosomes in the telomere. A Cdc13 defect leads to a defect in replication of one homolog. The defect leads to extensive ssDNA of the lagging strand template sister, and a complete leading strand template sister. The exposed lagging strand template may form a hairpin, allowing cleavage by Sae2 in the single copy sequence. The nicked lagging strand template is now the parental primer strand, which invades an intact homolog in single copy sequence. Two outcomes are possible: replication to the end of the chromosome or re-initiation of BIR. BIR may again abort, in which case the parental primer 3’ may be shortened, invade the homolog, and attempt replication again.

This model is essentially the Break Induced Replication (BIR) pathway, attractive to us given a link between telomeres and BIR (Özer & Hickson, 2018; Kramara *et al*, 2018; Anand *et al*, 2013). Determining whether failed BIR is indeed responsible for creating the initial unstable chromosome and its progression is a future research goal. Cycles of instability in sectored colonies bear some similarity to “cascades of instability” reported for a telomere-linked unrepairable DSB in a Chr III disome (Vasan *et al*, 2014), a system used to decipher molecular events of BIR. This model poses a plethora of predictions yet to be tested: the possible role of adaptation in cell cycle resuming with a linear unstable chromosome; the role of a hair pin in invasion; the role of the distance of *CAN1* from the telomere, etc.

### How might a telomeric unstable chromosome progress?

We suggest that an unstable chromosome forms near the telomere and from there progresses to assume one of many fates; recombination in various intervals, chromosome truncation at an internal TG sequence, dicentric formation via fusion of inverted repeats at the 403 site, or simple loss. We suggest these forms may arise through progressive degradation by Exo1 of a linear chromosome (Hackett & Greider, 2003). Given the “Failed BIR Model” in Figure 7, we suggest that at any point during degradation, strand invasion into the homolog may arise, resulting in a Class 1 recombinant. When degradation proceeds to the ^281^TG_126_ site, a truncated chromosome forms. When degradation proceeds to the 403 site, the inverted repeat sequences are exposed and an intra-sister fusion occurs, forming a dicentric. Extensive ssDNA may in fact be a feature of the unstable chromosome, perhaps even generating chromosomes “entangled” at their ends via extensive bonding between ssDNA regions. The notion of entangled chromosomes as a feature of the *cdc13*-induced unstable chromosomes is motivated by studies in fission yeast of a defect in *taz1^-^* cells; Taz1 protein facilitates DNA replication through the telomere, and in *taz1^-^*cells, entangled chromosomes occur, generating instability (Miller & Cooper, 2003). The nature of the telomere-proximal unstable chromosome in our studies may be, therefore, a form of dicentric after all, but not covalently bound.

## Materials and Methods

### Yeast Strains

All yeast strains used in this study were derived from the A364a TY200 disome strain previously described (Table S2; (Weinert & Hartwell, 1990; Admire *et al*, 2006; Paek *et al*, 2009)). The TY200 wild type Chr VII disome strain is *MATα +/hxk2::CAN1 lys5/+ cyh^r^/CYH^S^ trp5/+ leu1/+ cenVII ade6/+ +/ade3*, *ura3-52*. The endogenous *CAN1* gene on Chr V was mutated and inserted on one copy of Chr VII in place of *HXK2*. Additional strains were generated by LiAC/ssDNA/PEG transformation of TY200 with DNA fragments or plasmids. Strains were verified by genetic analysis and/or PCR. There were at least two separate strains made and analyzed for each mutant reported.

The various *cdc13* mutant strains was made by either Margherita Paschini or in house with a plasmid carrying the corresponding *cdc13* mutation via a “pop-in, pop-out” mechanism (pVL5439 (*cdc13^F684S^*); pVL7013 (*cdc13^Y556A,Y558A^*); gifts from V. Lundblad; (Paschini *et al*, 2012)). Double mutants with *cdc13^F684S^*were constructed by transforming PCR amplifications of *KanMX4*-marked genes replacements from the Gene Deletion Library into *cdc13^F684S^*strains using primers that flanked the coding region. It should be noted that one isolate of *cdc13^F684S^* has an altered Chr VIII and V. We do not believe that these alterations affect the frequency of chromosome instability since neither copy of Chr VII was affected and both variants of *cdc13^F684S^*, with and without the altered karyotype, had the same frequency of all three types of chromosome instability. Double mutants integrated into the *cdc13^F684S^*isolate with the altered karyotype are marked in the table in S2 Table.

The *URA3* proxy strains were generated as previously described (Paek *et al*, 2009). Briefly, the *URA3* gene fragments were linked to genes encoding drug resistance in two cassettes: RA:*NAT1*:RU and A3:*KanMX4*. These were inserted into plasmids containing ∼500 bp of sequence homology to flanking sites of the 403 kb (RA:*NAT1*:RU) or 535 kb (A3:*KanMX4*) regions of Chr VII. Plasmids were digested with the appropriate restriction enzymes to free the targeting fragment and *cdc13^F684S^*cells were transformed and selected for drug resistance. Correct insertions were verified by PCR and genetic analysis. In genetic analyses we verified that insertions were linked to *CAN1* (the changes were in the bottom homolog in Fig 1).

The ^281^TG:*URA3* strains were generated by PCR amplification of a 126 bp TG track from telomere IL from a pRS406:TG^126^ plasmid. The pRS406:TG^126^ plasmid was created by subcloning the TG fragment from a PCR 2.1 vector (gift from V. Lundblad) into pRS406 with restriction enzymes. The TG^126^ fragment from pRS406-TG^126^ was transformed into TY200 and *cdc13^F684S^*with primers containing 45 bp of homology to the DNA of the target sequence. *URA3* was amplified via PCR from the pRS406 plasmid. Cells were then transformed and selected for Ura^+^. Isolates were verified by PCR with primers outside of the targeted region. Further genetic analysis was performed to verify that insertions were integrated into the *CAN1* homologue of Chr VII.

The *^122^KanMX4* marker was integrated into *cdc13^F684S^* by transforming *cdc13^F684S^*with DNA fragments containing the *KanMX* gene flanked by 45 bp of homology to the Chr VII 122 kb locus. DNA fragments were synthesized by PCR amplification of the *KanMX4* gene from the pRS400 plasmid with primers containing 45 bp of homology to the target sequence. The *KanMX4* gene replaced Chr VII 121500 bp – 122100 bp. Correct insertion of the *KanMX4* construct was confirmed via PCR with primers outside of the targeted region, and the integrated of the construct into the *CAN1* homologue of Chr VII was confirmed by genetic analysis (Can^R^ colonies became sensitive to G418/geneticin).

The following drug concentrations were used whenever drug selection was appropriate: canavanine (Can; 60 μg/mL), G418/geneticin (100 μg/mL), hygromycin B (Hyg; 300 μg/mL), and nourseothricin (Nat; 50 μg/mL).

### Chromosome instability assays

Genetic analysis determined the frequency of unstable chromosomes, allelic recombinants, and chromosome loss by a slightly modified version of the previously described instability assay (Admire *et al*, 2006). Briefly, strains were struck to minimal media (Min + uracil) at 25 °C to select for retention of both Chr VII homologs. Single cells were then plated to rich media (YPD, 2% dextrose) and grown for 2-3 days at 30 °C to form colonies and allow instability to arise. Once colonies were formed, individual colonies were suspended in water, cell concentration was determined by counting cells with a haemocytometer, and a known number OD cells were plated to selective media and grown at 25 °C to measure chromosome instability.

To determine the frequency of chromosome loss cells were plated to selective media containing canavanine (60 μg/mL) and all essential amino acids except arginine and serine. Cells were grown at 25 °C for 5-6 days to allow for colony formation. Loss was determined by replica- plating to identify Ade^-^ colonies (Can^S^ Ade^-^). The frequency of chromosome loss was calculated by determining the ratio of the number of Can^R^ Ade^-^ colonies by the total number of cells originally plated.

To determine the frequency of unstable chromosomes and recombination, cells were plated to canavanine plates that also lacked adenine (in addition to arginine and serine). Cells were grown for 5-6 days at 25 °C and then colonies were scored based on morphology (sectored or round, Fig 1D). The frequency of sectored (unstable chromosomes) and round (recombinants) was calculated by determining the ratio of the number of Can^R^ Ade^+^ sectored or round colonies to the total number of cells originally plated. Averages and standard deviations were calculated from analysis of at least 6 colonies grown on rich media that were then plated to selective media. Unless otherwise stated, statistical tests were performed using the Mann-Whitney U method; P values are reported in Table S3.

### Phenotypic analysis of altered chromosomes

#### Single Colony Lineage Analysis

Cells from a single round or sectored Can^R^ Ade^+^ colony was isolated and suspended in water. Approximately 200 to 300 cells were then plated to rich media (YPD) and grown for 2-3 days at 25 °C to form colonies, each from a single cell. The phenotypes of each colony were determined by replica-plating to synthetic media, each lacking one essential amino acid, or media containing a drug (cycloheximide, 5 μg/ml). Cells were grown for 2-3 days at 25 °C, and growth on each selective plate was assessed. The proportion of Class 1 or 2 recombinants per region was calculated by dividing the number of genetic recombinants with the total number of recombinant colonies assayed. Colonies that lacked all the markers were scored separately as chromosome loss.

#### Pooled Colony Lineage Analysis

Cells from approximately 50 round or sectored Can^R^ Ade^+^ colonies were pooled and suspended in water. The retention and loss of individual genetic markers were analyzed as done in the Single Colony Lineage Analysis.

### Single cell cycle assay

Cells were grown to mid-log overnight at 25 °C in 100 ml of YPD. Cells were counted via hemocytometer to confirm that they were in mid-log (2-6×10^6^ cells/ml). Once the cells were at the appropriate concentration, hydroxyurea was added to 20 ml of culture (0.2 M HU) and the culture was incubated at 25 °C for 3 hours. Cells were then washed twice with H_2_O and resuspended in 20 ml of YPD (t_0_). The culture was then split; half of the culture was incubated at 37 °C while the other half remained at 25 °C. The cultures were incubated at either 37 °C or 25 °C for a total of 4 hours. To measure chromosome instability, 1 ml of culture was taken from both cultures every hour after the temperature shift, as well as from the t_0_ time point. The 1 mL aliquots were sonicated, counted, and plated to CAN and CAN-ade for the instability assay within 30 minutes of collection (instability assay described above). Cells were also plated to selective plates lacking arginine to measure cell viability. Viability was measured after 1 day of growth by counting the number of viable microcolonies versus single cells. For the control, cells were never exposed to HU and were instead shifted to either 37 °C or 25 °C immediately after they were confirmed to be in mid-log.

Samples for FACS were taken at mid-log, after HU was washed out (if applicable), and then at 0:15, 0:30, 0:45, 1:00, 2:00, 3:00, and 4:00 hours after cells were shifted to either 37 °C or 25 °C. The section below describes the FACS protocol.

To inactivate *cdc13^F684S^* during S phase, cells were grown to mid-log overnight in 100 ml of YPD at 25 °C. Once in mid-log growth, a 1 ml aliquot was taken for the instability assay and 10 ml of culture was incubated at 37 °C. After 4 hours at 37 °C another 1 ml aliquot of cells was taken for the instability assay.

To limit *cdc13^F684S^* inactivation to G2, cells were grown to mid-log overnight in 100 ml of YPD at 25 °C. Once in mid-log, nocodazole was added to 20 ml of culture (10 μg/ml, 1% DMSO) and the culture was incubated at 25 °C for 3 hours. After incubation, cells were washed twice with water and resuspended in 20 mL of fresh YPD (t_0_). The culture was then incubated at 37 °C for 4 hours. Aliquots of cells were taken at t_0_ and after 4 hours at 37 °C for the instability assay. To hold cells in G2/M after the temperature shift additional nocodazole was added (10 μg/ml; 1% DMSO).

As done previously, the 1 mL aliquots for the instability assay were sonicated, counted, and plated to CAN and CAN-ade for the instable assay within 30 minutes of collection. Cells were also plated to selective plates lacking arginine to measure cell viability.

Samples of FACS analysis were taken at mid-log, after noc was washed out (if applicable), and at 0:30, 1:00, 2:00, 3:00, and 4:00 after the cells were shifted to 37 °C. The FACS protocol is described in the following section.

### FACS analysis

Cells were pelleted from 1 mL of log YPD culture and fixed in 70% ethanol overnight at 4 °C. Ethanol was removed and cells were resuspended in 50 μl of 50 mM sodium citrate (pH 7.4) and sonicated at low power (8 s at 20% power). Cells were pelleted again and resuspended in 1.0 ml of 50 mM sodium citrate with 0.25 mg/ml of RNaseA for 1 hour at 50 °C. 25 μl of proteinase K solution (20 mg/ml proteinase K in 10 mM Tris pH 8.0, 1 mM CaCl_2_, 50% glycerol) was subsequently added and the cells were incubated at 50 °C for an additional hour. Cells were then pelleted and resuspended in 1.0 ml of 50 mM sodium citrate containing 2 µM Sytox green (Invitrogen). Cells were left to incorporate the dye overnight at 4 °C. Samples were scanned using a Attune Acoustic Focusing Flow Cytometer (Invitrogen).

### Pulsefield gel electrophoresis and Southern analysis

To identify altered chromosomes, pulsefield gel electrophoresis and Southern blotting for chromosome VII were performed as described (Admire *et al*, 2006). Southern hybridization was performed using a P^32^-labeled probe to Chr VII sequences 503875 bp–505092 bp as previously described (Beyer & Weinert, 2016).

### In-gel hybridization

750 ng – 1 µg of XhoI digested genomic DNA was subjected to agarose gel electrophoresis (0.7% agarose) for 16 hours at 60 V. The gel was placed on a double-layer of Whatman paper with a piece of plastic wrap over top and mounted on a Bio-rad (model 583) gel drier. Drying was carried out for ∼20 minutes at room temperature. The dried gel was sealed in a plastic bag and hybridized with minimum 1 million CPMs of γ-ATP32 end-labeled oligonucleotide 5’-CCCACCCACCACACACACCCACACCC-3’ in in-gel hybridization buffer overnight in a 37°C water bath. After removal of excess hybridization buffer, the gel was washed 3-4 times for 30 minutes in 0.25X SSC at room temperature with agitation, sealed in a bag, and exposed to Amersham Hyperfilm MS or phosphor cassette. After, the in-gel was subjected to a denaturing Southern blot in order to visualize the total telomeric DNA. The Southern blot was performed as follows: 10 minutes in 0.25M HCl, 45 minutes in Southern denaturing solution (1.5M NaCl, 0.5M NaOH), 5 minutes 0.4M NaOH. It was then transferred to a Hybond-XL nylon membrane. After transfer, the membrane was prehybridized for 1 hour in Church buffer and then hybridized overnight at 65°C with a radiolabeled probe that hybridizes to telomeric repeats. The membrane was washed for 20 minutes at room temperature in 2X SSC and exposed on a phosphor cassette.

## Acknowledgments

We thank Vicki Lundblad and Margherita Paschini for the *cdc13^F684S^* and *cdc13^Y556A,Y558A^*mutants in addition to helpful discussions, plasmids, and construction of strains. We thank Shang Li for plasmids. We also thank Lisa Shanks, Tracey Beyer, and Peter Vinton for frequent discussions of this work. T.W. and R.E.L were funded through the NIH GM076186-5 and training grant GM08659 to R.E.L.

## Competing Interests

We declare that no financial or non-financial competing interests exist.

---

**Fig S1.**
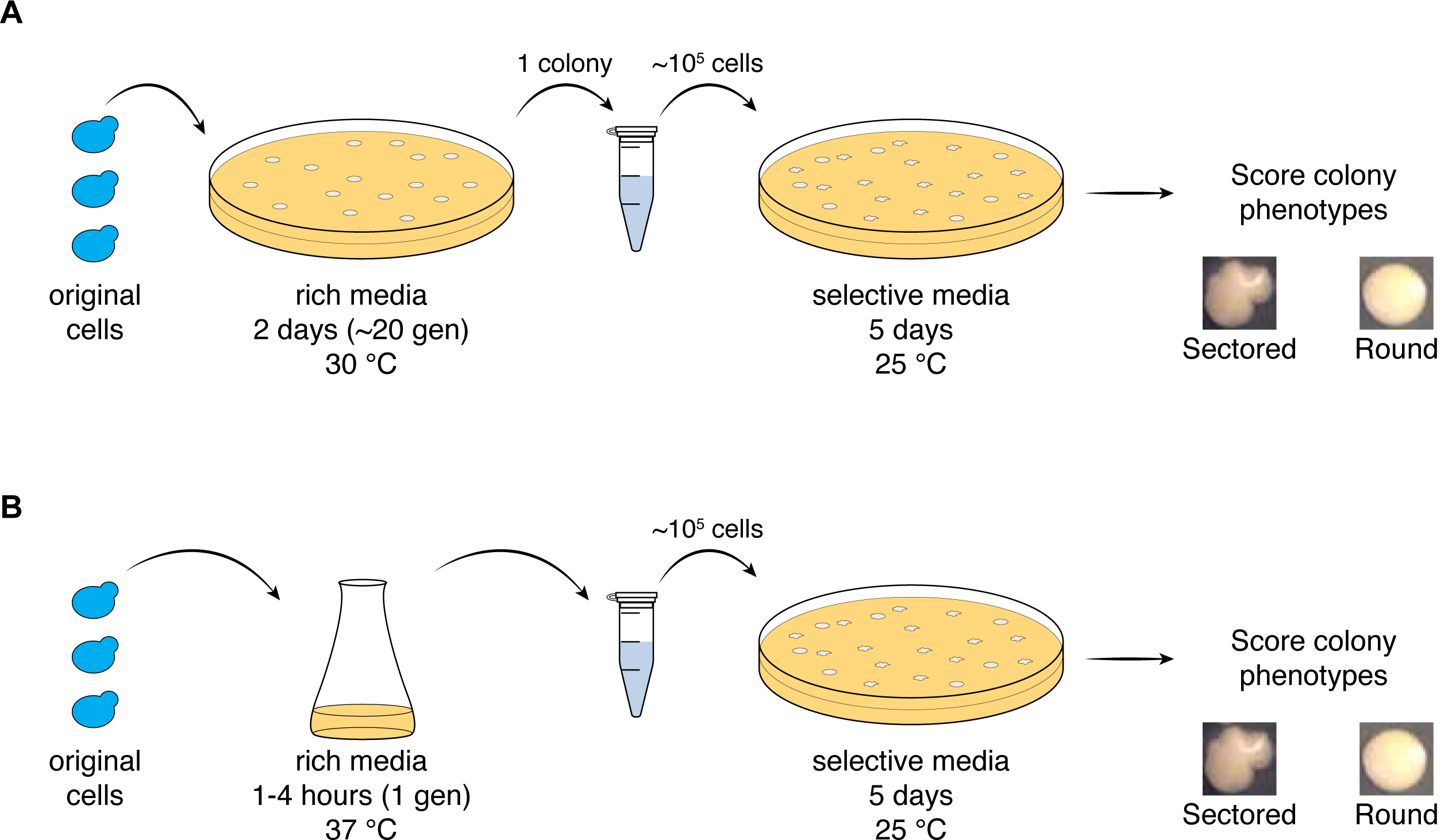
Diagram of the instability assay. **(A)** Cells are plated to rich media for 2 days at 30 °C (semi-restrictive temperature for *cdc13^F684S^*), then individual colonies are plated to canavanine-containing media for 5 days at 25 °C to select for chromosomal rearrangements. Instability is scored by comparing the number of “sectored” or “round” colonies vs the number of cells originally plated. **(B)** Asynchronous cells are grown at 25 °C then shifted to 37 °C for 1-4h. Then ∼10^5^ cells are plated to canavanine-containing media for 5 days at 25 °C to select for chromosomal rearrangements. Instability is scored by comparing the number of “sectored” or “round” colonies vs the number of cells originally plated.

**Fig S2.**
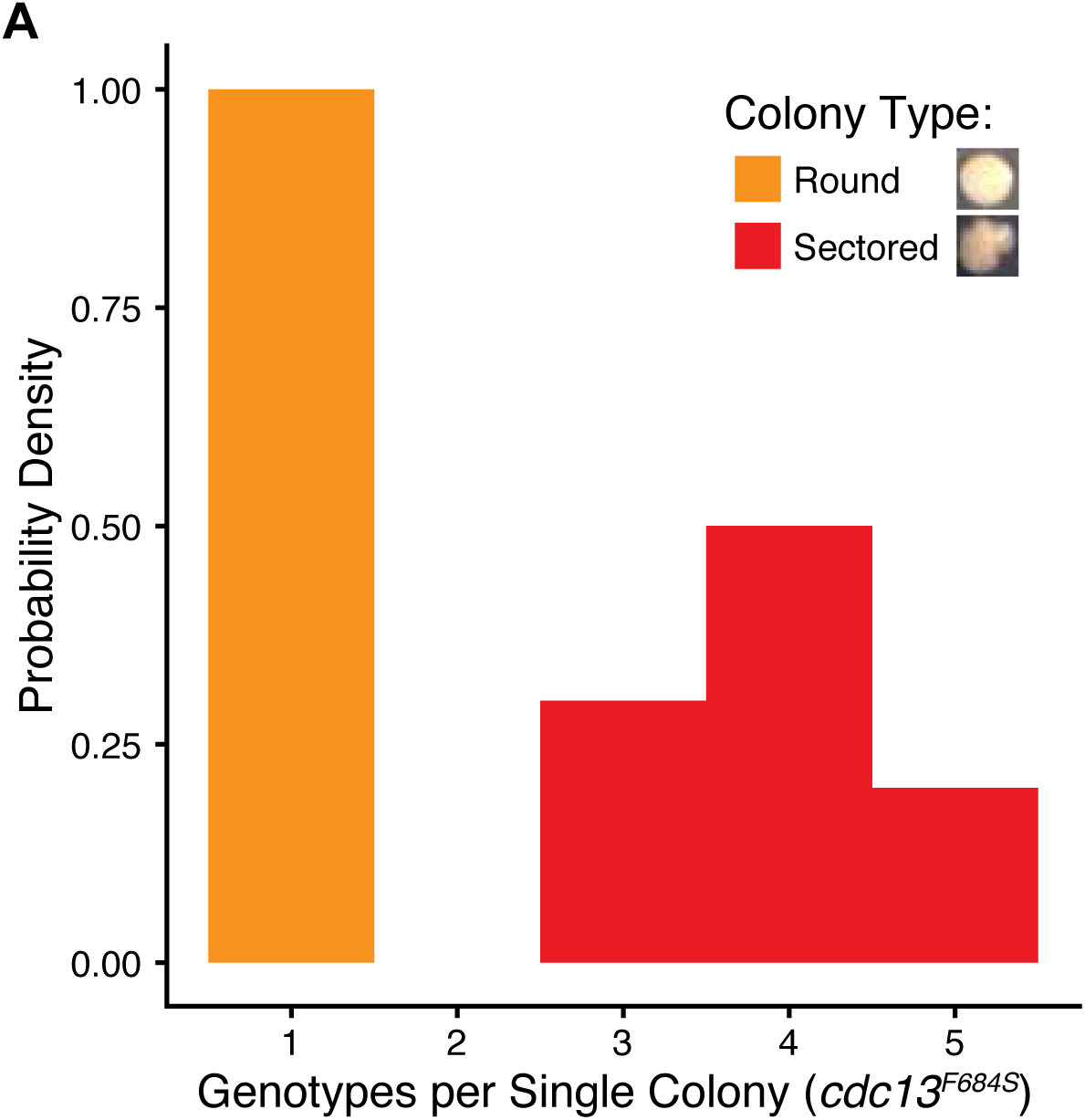
Sectored colonies contain multiple genotypes. **(A)** Histogram for the number of genotypes in sectored (red) and round (orange) colonies, n = 20 (n = 10 colonies each for sectored and round).

**Fig S3.**
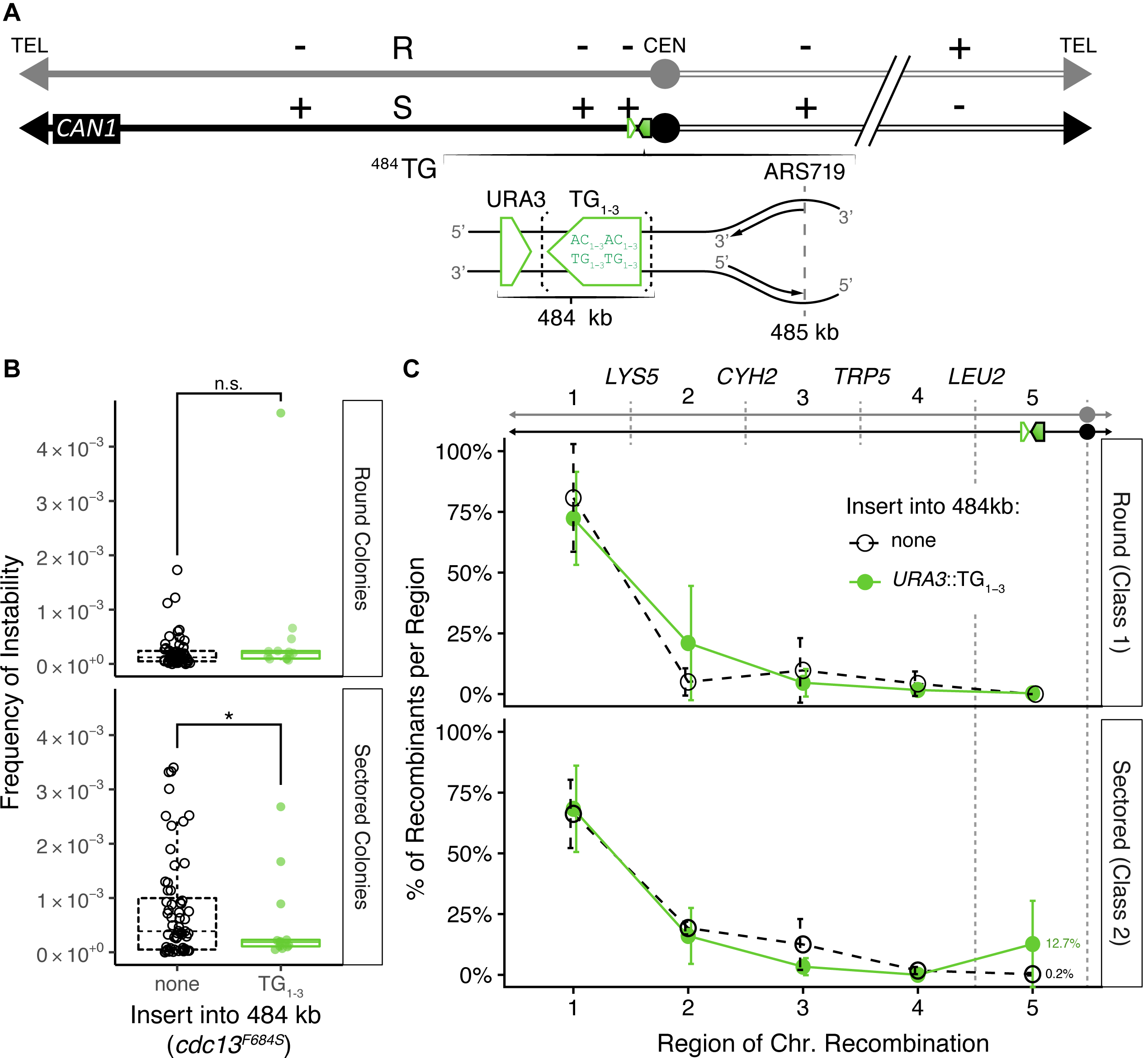
Inserting the TG_1-3_ repeat into the 484 kb locus does not change the distribution of Class 1 recombinants in round colonies, but does change in distribution in sectored colonies. **(A)** Diagram for the TG repeat (green box) is insertion in the 484 kb locus. **(B)** Frequency of round (*top*) and sectored colonies (*bottom*) from an unmodified *cdc13^F684S^* and *cdc13^F684S^ ^484^TG_1-3_* are not altered. The mean and standard deviation are shown (n > 12; n.s. > 0.05; Mann-Whitney U). **(C)** Distribution of genotypes from round (Class 1 recombinants) and sectored colonies (Class 2 recombinants) from *cdc13^F684S^* with no insert or with the TG_1-3_ insert. The average percentage and standard deviation for 3 independent experiments are shown.

**Fig S4.**
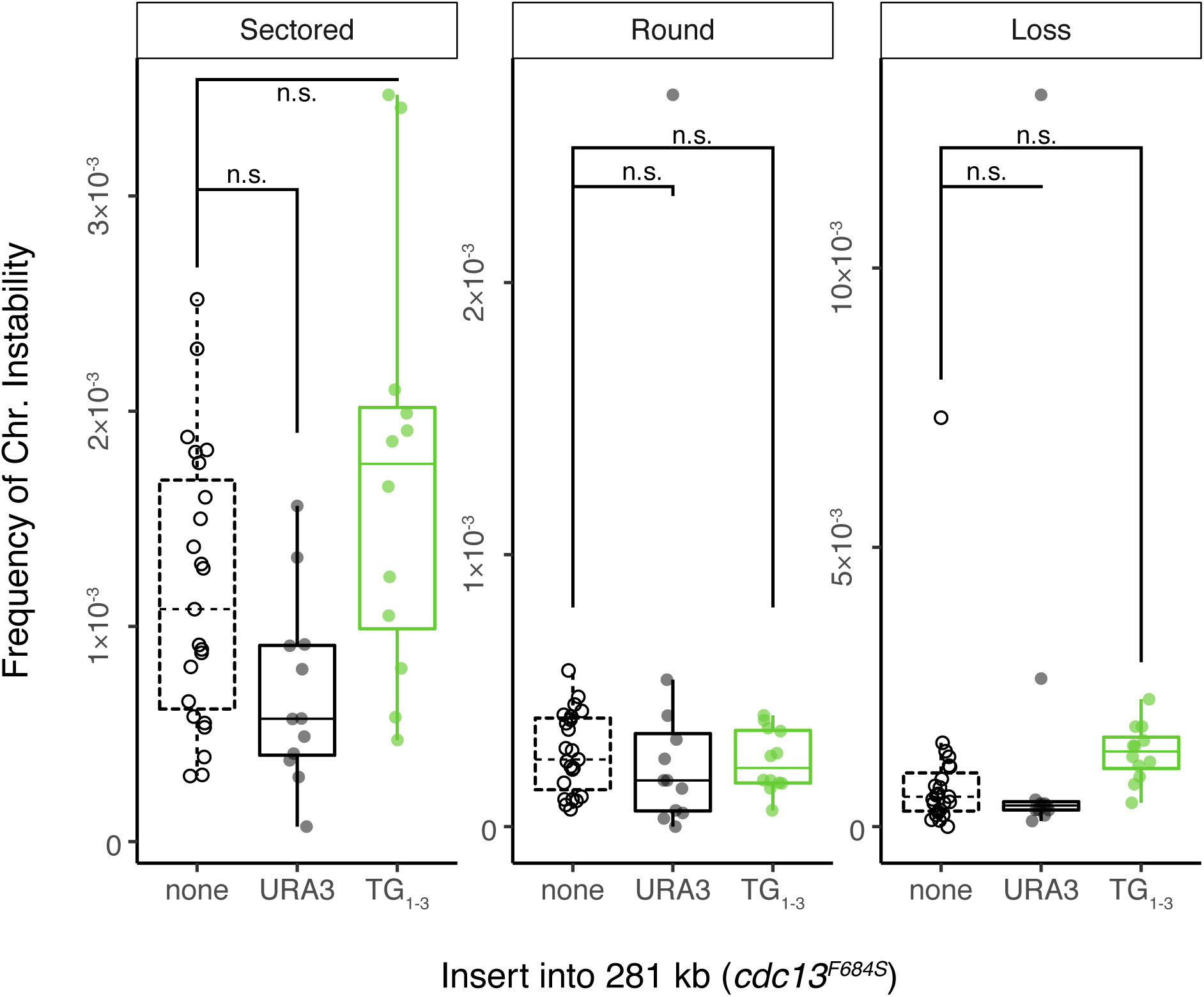
Inserting the TG_1-3_ repeat into the 281 kb locus does not alter the frequency of unstable chromosomes, recombination, or chromosome loss. **(A)** Frequency of instability for sectored, round, and loss colonies from an unmodified *cdc13^F684S^*, *cdc13^F684S^ URA3*, and *cdc13^F684S^ ^281^TG_1-3_*. The mean and standard deviation are shown (n > 12; n.s. > 0.05; Mann-Whitney U).

**Fig S5.**
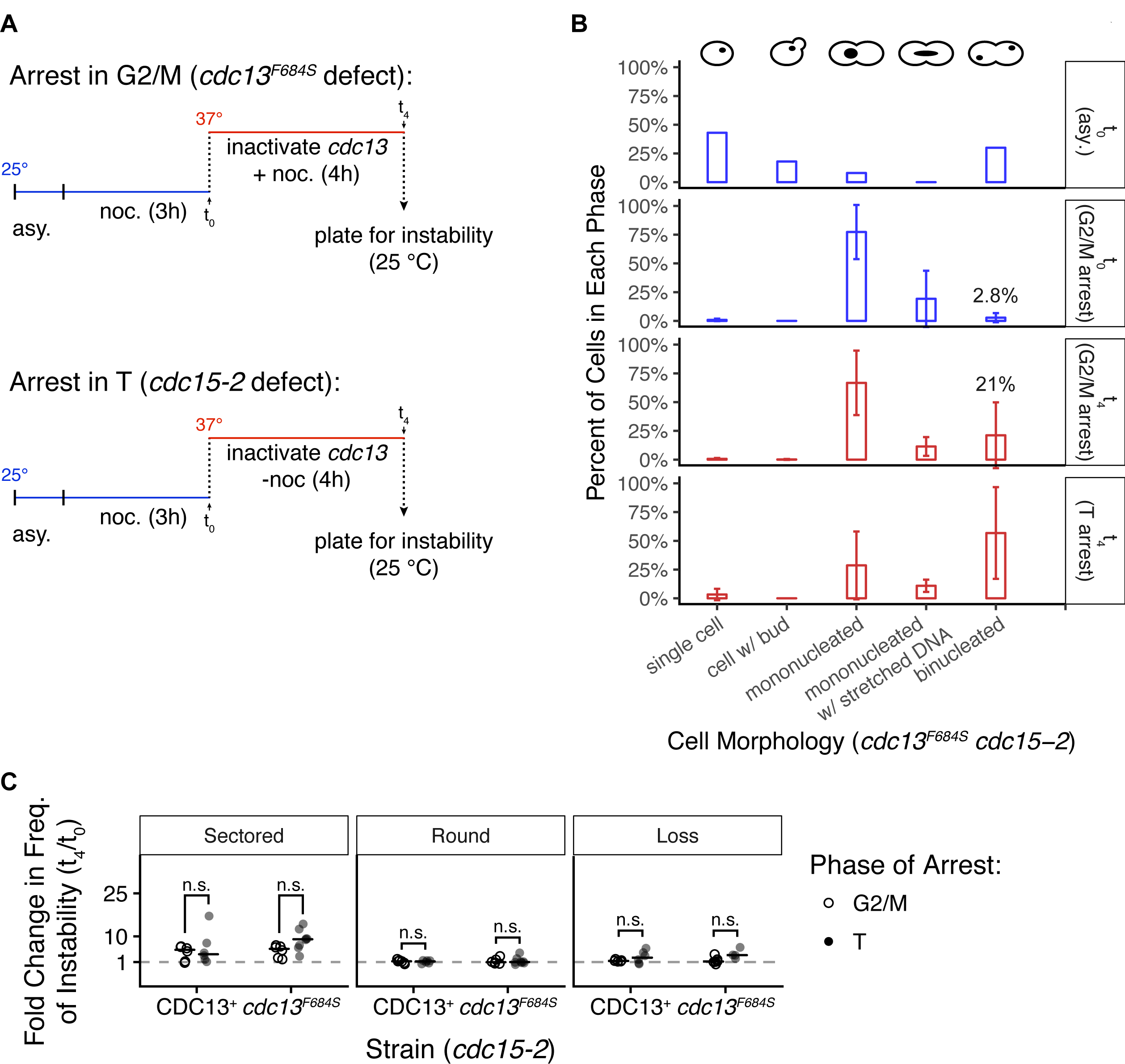
cdc13^F684S^ activity in S and inactivity for either only G2 or only G2-T does not result in unstable chromosomes. **(A)** Schematic for arresting cells in either G2/M or T phase. Cells were grown at 25 °C until they reached mid-log (t_asy_), then were incubated with nocodazole for 3 hours. Cells were then washed (t_0_) and the culture was split. Both halves were incubated at 37 °C for 4 hours, one with (t_4_ + noc; *left*) and without (t_4_ – noc; *right*). At t_asy_, t_0_, t_4_ + noc, and t_4_ - noc cells were collected for DAPI staining and the instability assay. **(B)** Nuclear morphology based on DAPI staining of chromosomes. The proportion of *cdc13^F684S^ cdc15-2* cells with each morphology for the different arresting protocols is shown. Blue: cells grown at 25 °C; red: cells grown at 37 °C. Mean and standard deviation from at least 6 independent experiments are shown. **(C)** Fold change in the frequency of sectored, round, and loss colonies (t_4_/t_0_) from *cdc15-2* CDC13^+^ and *cdc13^F684S^ cdc15-2* with (n = 6; 6) and without (n = 6; 8) additional nocodazole added after the initial noc arrest. The fold change between t_4_ and t_0_ remained low despite the arresting conditions (* < 0.05, Mann-Whitney U).

**Fig S6.**
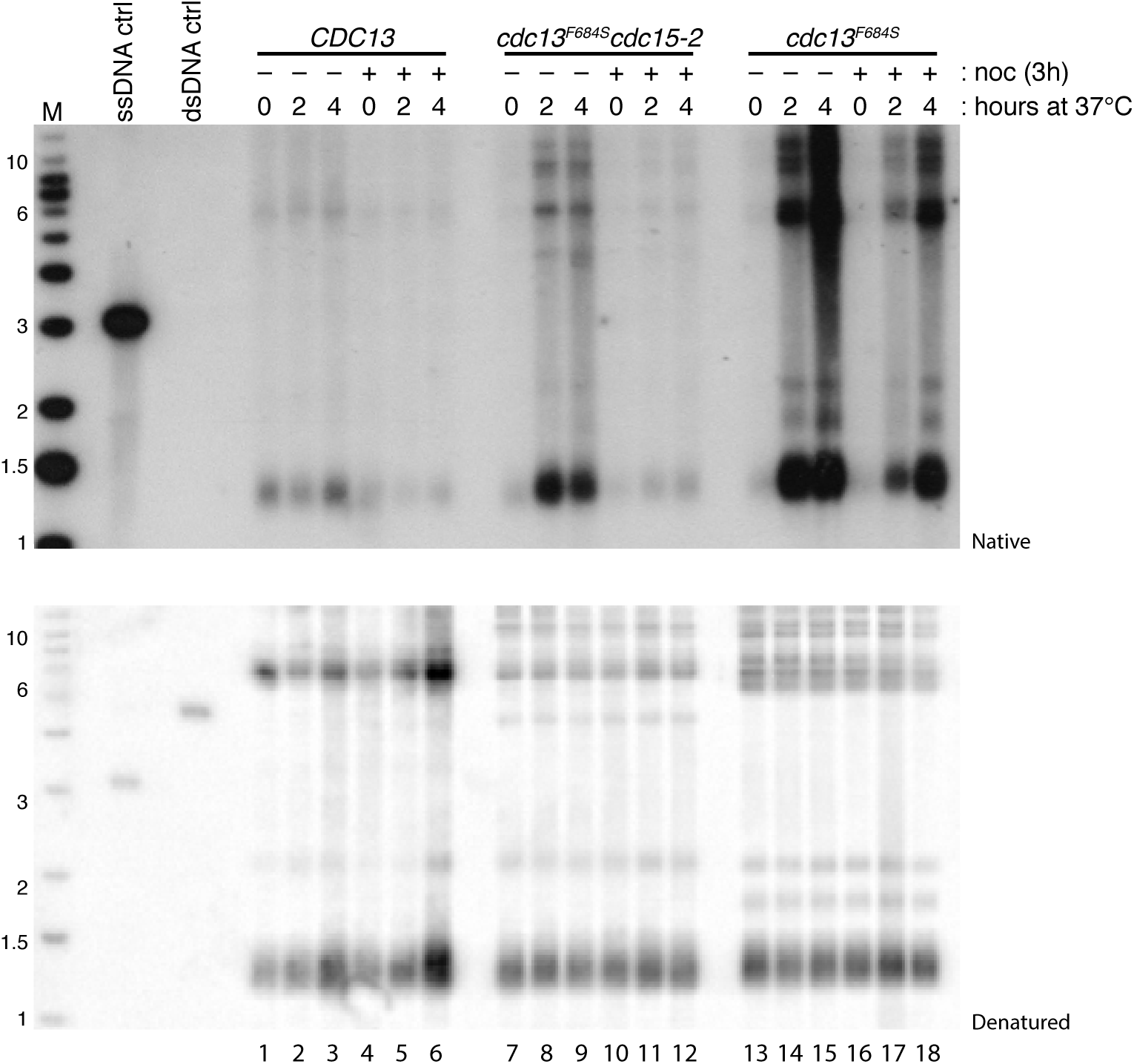
ssDNA generated in cdc13^F684S^ during G2/M probably comes from escaped cells in the next cell cycle. Uncropped image of the non-denaturing in-gel hybridization in 3E. Note that ssDNA is generated in *cdc13^F684S^* after noc arrest at 25 °C and release into 37 °C (lanes 13-18; not depicted in 3E).

**Fig S7.**
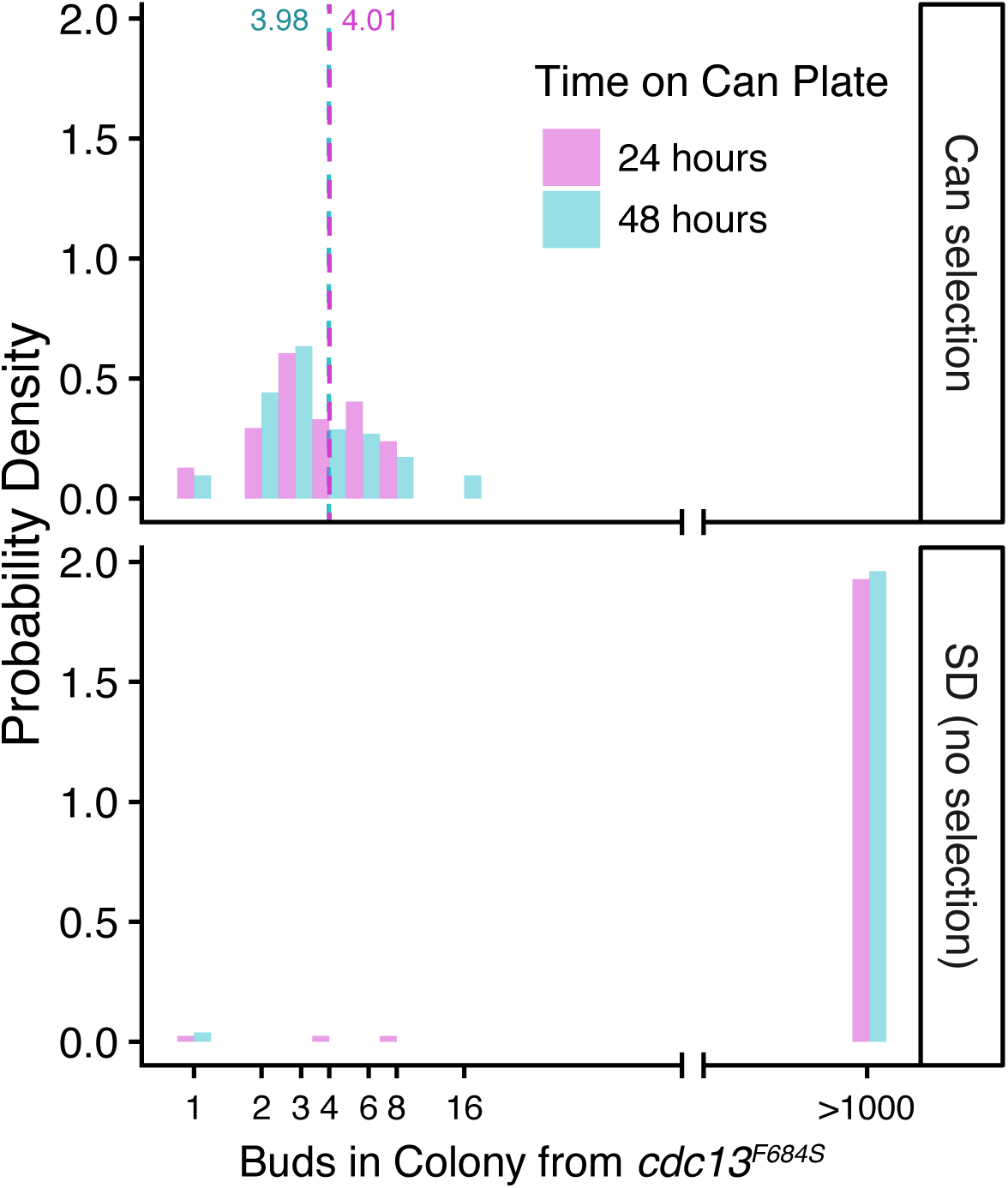
cdc13^F684S^ cells proceed through 1-3 cell divisions before dying on canavanine. Density curve for the colony size of *cdc13^F684S^* (t_0_ cells from Figure 3A *top* grown on plates with (*top*) or without canavanine (*bottom*). Colony size was scored at 24h (purple) and 48h (blue). Dashed red line: the average number of buds per colony for cells grown with canavanine (mean = 4.01 buds).

**Fig S8.**
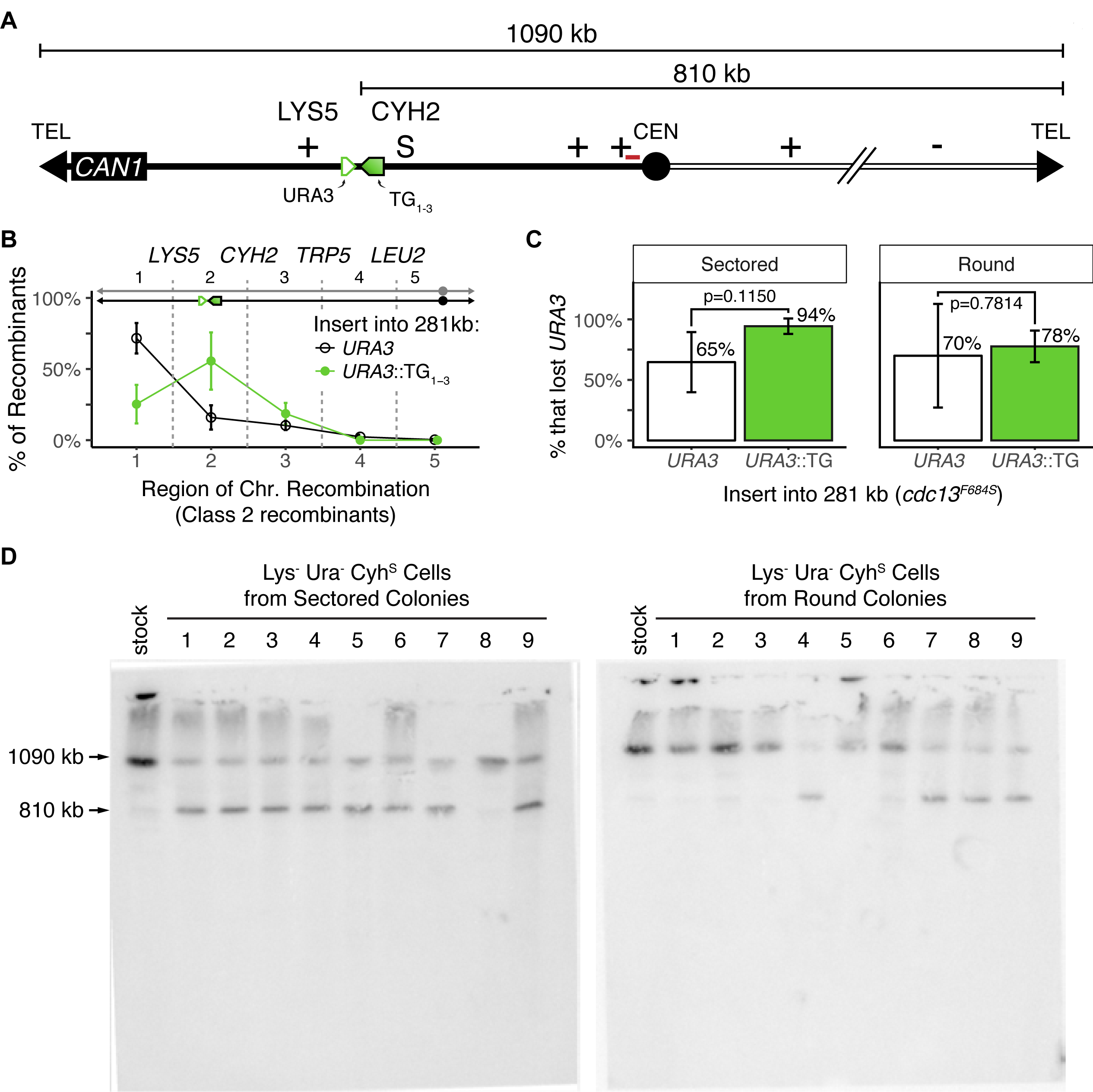
Unstable chromosomes can resolve by forming truncations at an internal TG repeat. **(A)** The ^281^URA3 or ^281^TG repeat (green box) inserted between *LYS5* and *CYH2* as in Fig 2B. Red line indicates the probe binding site. **(B)** Distribution of Class 2 recombinants from sectored colonies from *cdc13^F684S^*with *URA3* or *URA3*::TG_1-3_ inserted into the 281 kb locus. The average percentage and standard deviation for 3 independent experiments are shown. **(C)** Proportion of Lys^-^ Cyh^S^ cells that have additionally lost *URA3* from sectored and round colonies. P values were calculate with a *t* test. **(D)** Southern blot of pulse-field gels from Lys^-^ Ura^-^ Cyh^S^ cells from sectored or round colonies. A Chr VII centromere-linked probe was used to label chromosome VII. The upper band corresponds to the normal Chr VII size (1090 kb) while the lower band corresponds to a truncation at 281 kb (810 kb; localizes with Chr II, 813 kb).

**Fig S9.**
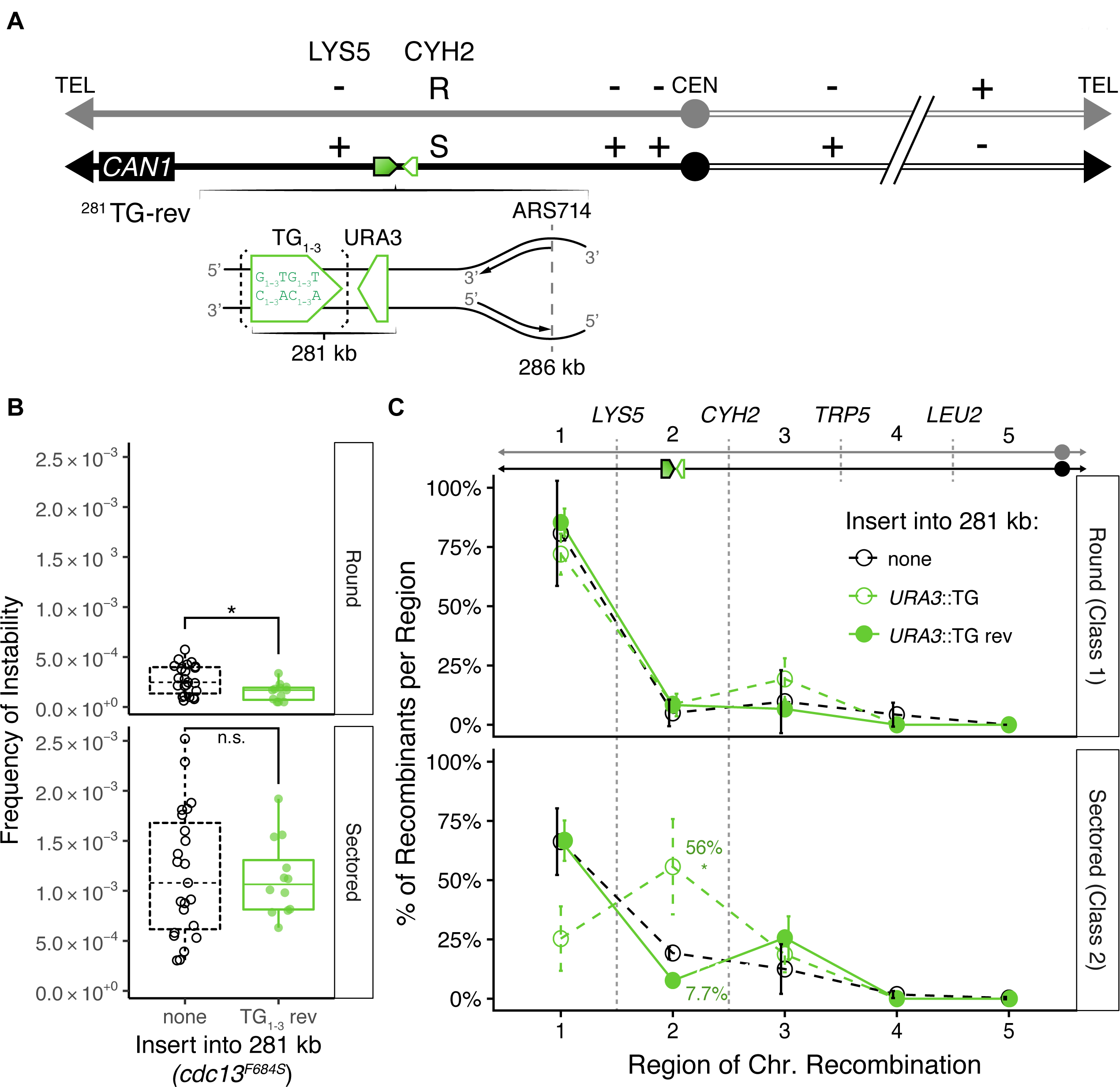
Inserting the TG_1-3_ repeat in the reversed orientation does not result in either an increase in instability or in a “spike” in TG-associated Class 1 or Class 2 recombinants. **(A)** The TG repeat (green box) in the 281 kb in the reversed orientation (TG-rich strand acting as the template for leading strand replication). **(B)** Frequency of round (*top*) and sectored colonies (*bottom*) from an unmodified *cdc13^F684S^* (white) and *cdc13^F684S^ ^281^TG_1-3_-*rev (green) are not altered. The mean and standard deviation are shown (n > 12; n.s. > 0.05; Mann-Whitney U). **(C)** Distribution of genotypes from round (Class 1 recombinants) and sectored colonies (Class 2 recombinants) from *cdc13^F684S^* with no insert, with ^281^TG_1-3_, or with ^281^TG_1-3_-rev. The average percentage and standard deviation for 3 independent experiments are shown (P = 0.01; one sample *t* test)

**S1 Table.**
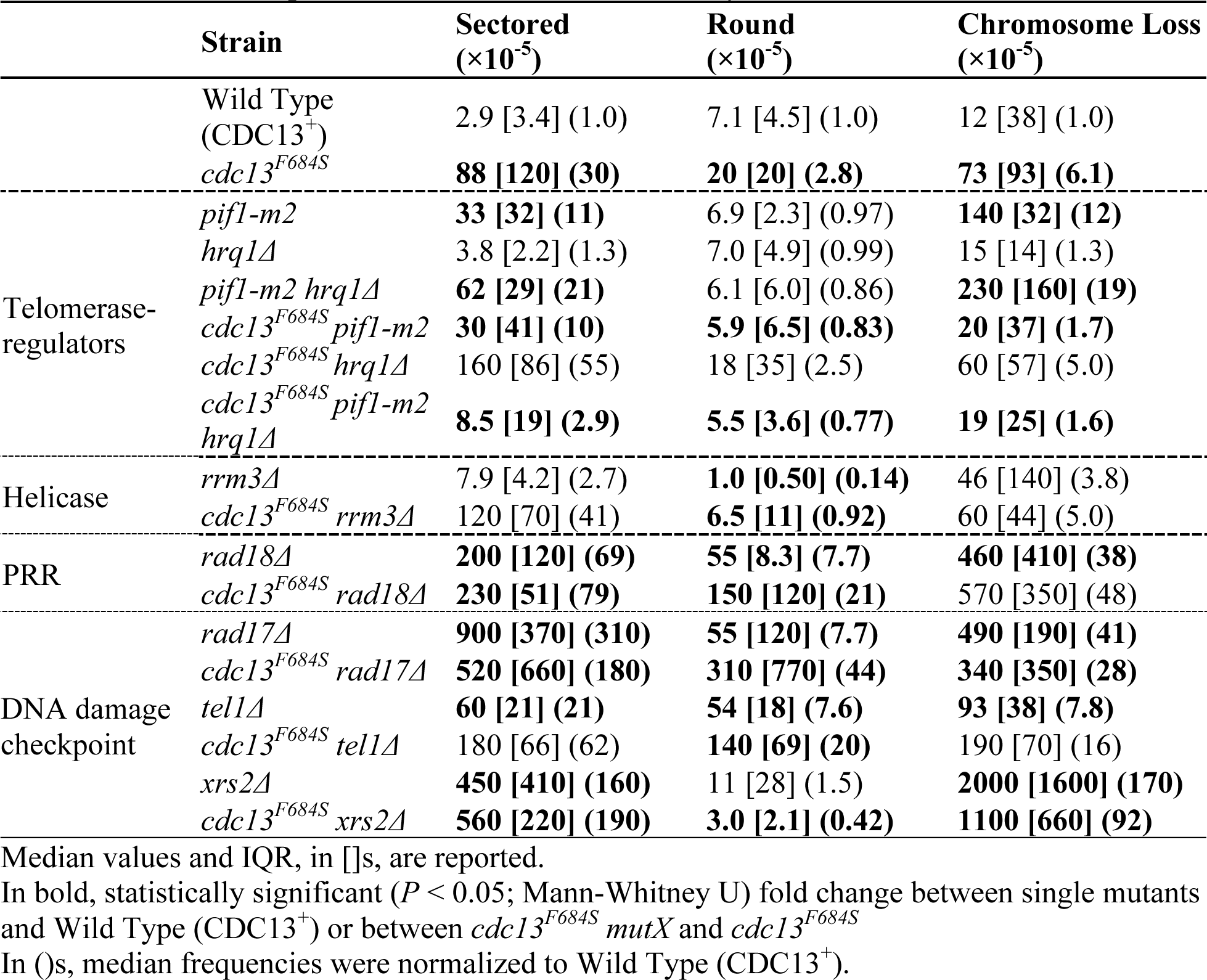
Median frequencies of chromosome instability in additional *cdc13^F684S^* mutants

**S2 Table.**
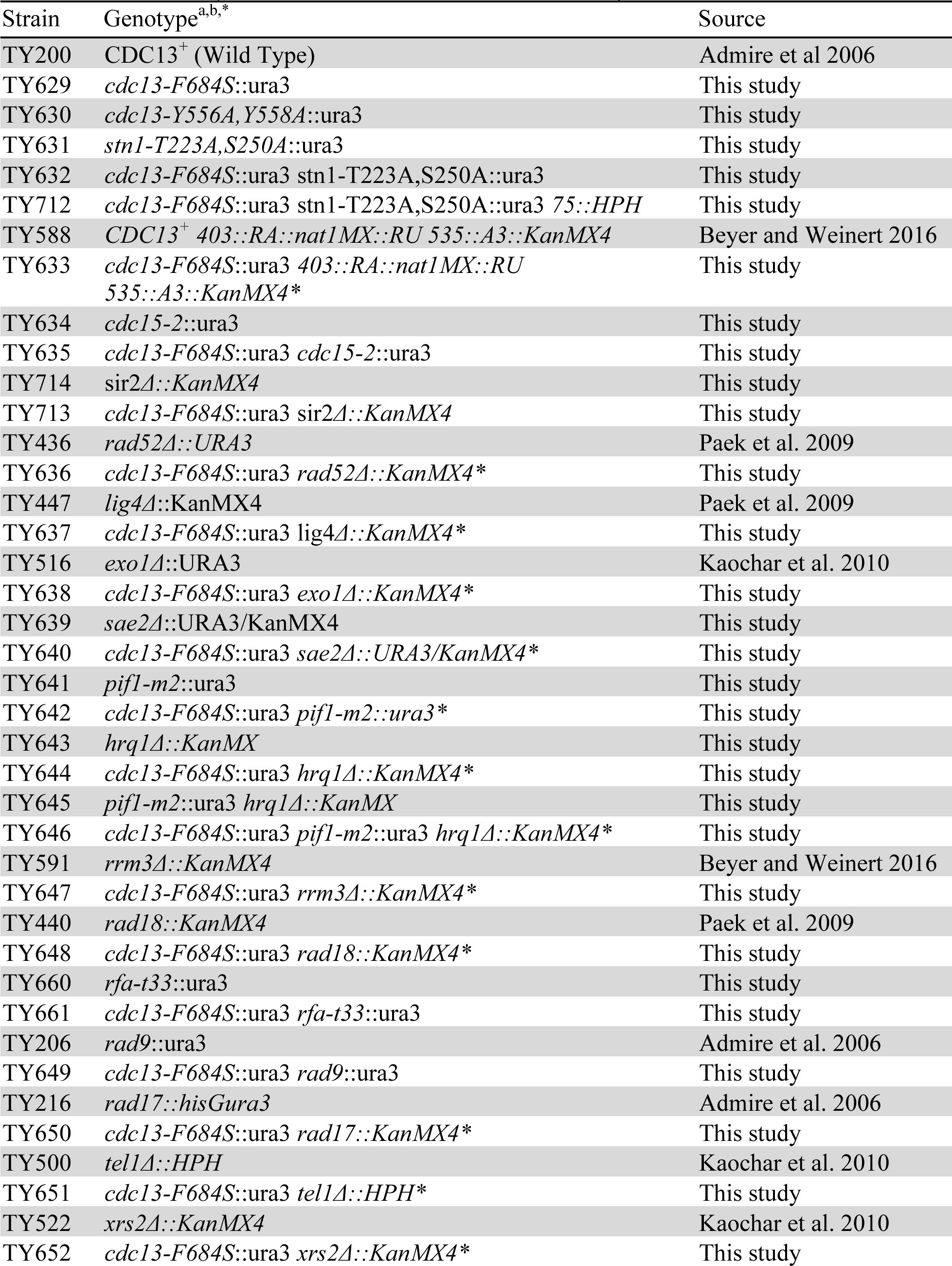
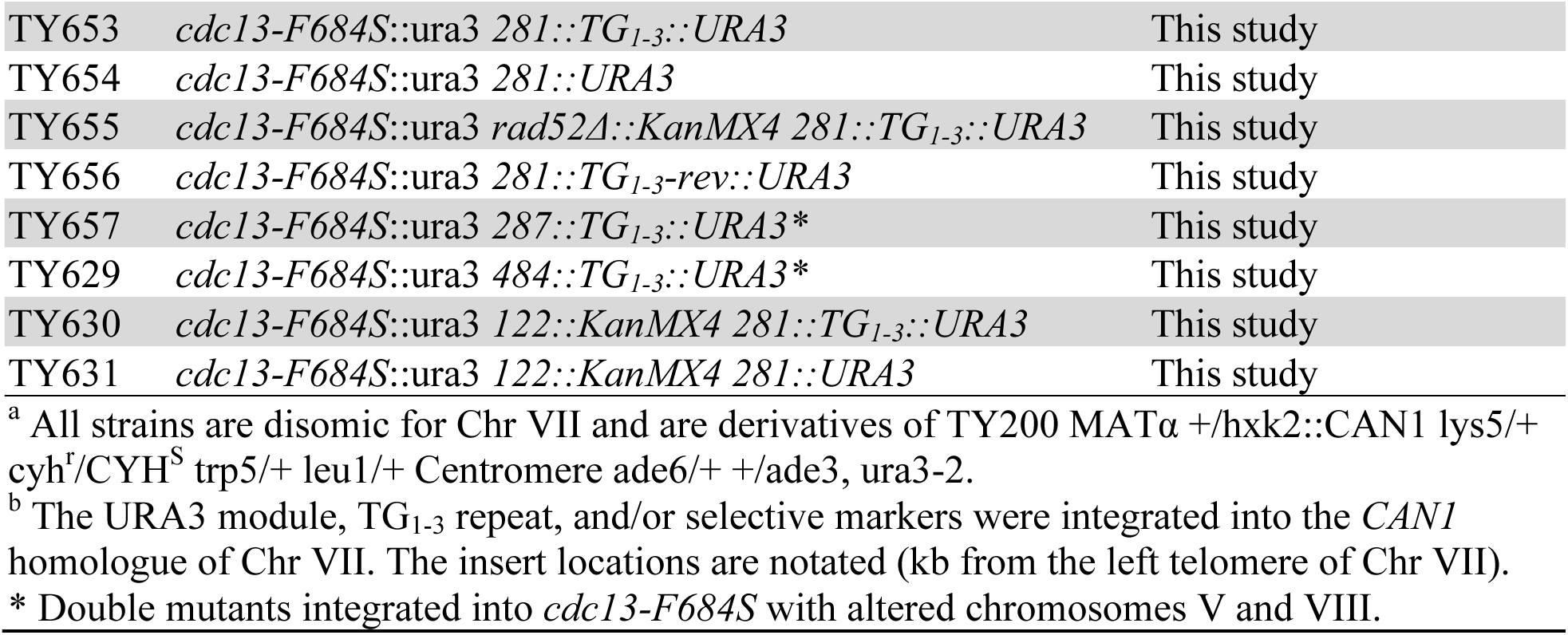
Saccharomyces cerevisiae strains used in this study

**Table S3.** List of P values

| P values | Sectored |  |  | Round |  |  | Loss |  |  |
| --- | --- | --- | --- | --- | --- | --- | --- | --- | --- |
| Strain | WT | *cdc13<sup>F684S</sup>* | other | WT | *cdc13<sup>F684S</sup>* | other | WT | *cdc13<sup>F684S</sup>* | other |
| Wild Type (CDC13<sup>+</sup>) | 1 |  |  | 1 |  |  | 1 |  |  |
| *cdc13<sup>F684S</sup>* | 2,15E-15 |  |  | 1,19E-07 |  |  | 3,83E-06 |  |  |
| *cdc13<sup>Y556A,Y558A</sup>* | 4,52E-08 |  |  | 0,000368 |  |  | 3,71E-07 |  |  |
| *stn1<sup>Y223A,S250A</sup>* | 0,7232 |  |  | 0,8115 |  |  | 0,6102 |  |  |
| *cdc13<sup>F684S</sup> stn1<sup>Y223A,S250A</sup>* | 6,11E-09 | 3,86E-06 | 3,20E-07 | 1,30E-07 | 0,1831 | 0,0002409 | 3,67E-07 | 0,002677 | 7,60E-07 |
| *rad52Δ* | 0,000129 |  |  | 0,000215 |  |  | 0,8246 |  |  |
| *cdc13<sup>F684S</sup> rad52Δ* | 1,16E-08 | 0,03403 | 0,003657 | 1,73E-08 | 2,37E-10 | NA | 9,54E-07 | 1,01E-07 | 0,0001616 |
| *lig4Δ* | 0,8226 |  |  | 0,473 |  |  | 0,5777 |  |  |
| *cdc13<sup>F684S</sup> lig4Δ* | 0,001286 | 0,3853 | 0,006812 | 0,2228 | 0,463 | 0,4021 | 0,5045 | 0,0006148 | 0,8838 |
| *exo1Δ* | 0,000981 |  |  | 3,15E-06 |  |  | 0,01449 |  |  |
| *cdc13<sup>F684S</sup> exo1Δ* | 0,000488 | 3,36E-11 | 0,1489 | 7,91E-10 | 2,49E-12 | 6,76E-05 | 0,5111 | 4,57E-07 | 0,001025 |
| *sae2Δ* | 0,004162 |  |  | 0,06667 |  |  | 0,1779 |  |  |
| *cdc13<sup>F684S</sup> sae2Δ* | 4,41E-07 | 0,0001274 | 0,000879 | 0,08548 | 0,01576 | 0,4371 | 0,0002404 | 0,4085 | 0,01188 |
| *pif1-m2* | 0,00013 |  |  | 0,7261 |  |  | 0,001046 |  |  |
| *hrq1Δ* | 0,8535 |  |  | 0,818 |  |  | 0,6248 |  |  |
| *pif1-m2 hrq1Δ* | 1,01E-06 |  |  | 0,3029 |  |  | 5,46E-05 |  |  |
| *cdc13<sup>F684S</sup> pif1-m2* | 3,53E-06 | 1,25E-06 | 0,5953 | 0,1717 | 1,27E-06 | 0,5679 | 0,9604 | 1,09E-06 | 0,0008232 |
| *cdc13<sup>F684S</sup> hrq1Δ* | 4,41E-07 | 0,01651 | 3,63E-05 | 0,005507 | 0,9464 | 0,4455 | 0,005741 | 0,4455 | 0,0005246 |
| *cdc13<sup>F684S</sup> pif1-m2 hrq1Δ* | 1,18E-05 | 5,77E-08 | 0,00069 | 0,07467 | 2,18E-07 | 0,9433 | 0,9801 | 1,68E-06 | 4,70E-05 |
| *rrm3Δ* | 0,01783 |  |  | 0,009511 |  |  | 0,09998 |  |  |
| *cdc13<sup>F684S</sup> rrm3Δ* | 2,12E-10 | 0,222 | 0,00181 | 0,6651 | 0,0007763 | 0,002507 | 0,0004666 | 0,2778 | 0,8181 |
| *rad18Δ* | 0,00013 |  |  | 0,0001627 |  |  | 0,009482 |  |  |
| *cdc13<sup>F684S</sup> rad18Δ* | 1,01E-06 | 0,0001672 | 0,09825 | 1,15E-06 | 2,92E-07 | 0,001131 | 1,27E-06 | 2,16E-06 | 0,3011 |
| *rfa1-t33* | 4,41E-07 |  |  | 0,002213 |  |  | 4,16E-07 |  |  |
| *cdc13<sup>F684S</sup> rfa1-t33* | 4,41E-07 | 1,04E-05 | 0,6297 | 0,01916 | 0,01676 | 0,05186 | 4,77E-07 | 1,79E-07 | 0,000656 |
| *rfa1-t33 sae2Δ* | 4,41E-07 | 1,48E-06 | 0,000108 | 0,03133 | 0,0008579 | 0,002047 | 4,16E-07 | 0,000656 | 0,0001077 |
| *rad9Δ* | 2,13E-10 |  |  | 0,5134 |  |  | 0,5212 |  |  |
| *cdc13<sup>F684S</sup> rad9Δ* | 5,13E-11 | 2,70E-07 | 4,32E-06 | 0,0007162 | 0,02695 | 0,01112 | 6,68E-11 | 1,10E-12 | 8,16E-08 |
| *rad17Δ* | 2,43E-06 |  |  | 0,001855 |  |  | 4,88E-05 |  |  |
| *cdc13<sup>F684S</sup> rad17Δ* | 1,16E-08 | 5,57E-07 | 0,05204 | 5,36E-07 | 2,06E-08 | 0,008988 | 6,98E-07 | 1,10E-05 | 0,2529 |
| *tel1Δ* | 0,00013 |  |  | 0,0001378 |  |  | 0,009482 |  |  |
| *cdc13<sup>F684S</sup> tel1Δ* | 0,000129 | 0,03207 | 0,004998 | 0,0001377 | 9,51E-05 | 0,06508 | 0,0007842 | 0,03053 | 0,02597 |
| *xrs2Δ* | 0,00013 |  |  | 0,2401 |  |  | 0,0001226 |  |  |
| *cdc13<sup>F684S</sup> xrs2Δ* | 1,01E-06 | 2,13E-07 | 0,6151 | 0,0009038 | 2,54E-06 | 0,03652 | 9,54E-07 | 3,56E-07 | 0,2475 |

*cdc13* variantsUnmodified *cdc13-F684S*
| Insertion into *cdc13-F684S* | Sectored | Allelic | Loss |
| --- | --- | --- | --- |
| 281 TG | 0,1148 | 0,5316 | 0,001654 |
| 281 *URA3* | 0,03112 | 0,3219 | 0,3292 |
| 281 TG rev | 0,9863 | 0,0136 | 0,8974 |
| 484 TG | 0,005027 | 0,5365 | 0,3255 |
| *URA3* proxy | 0,01332 | 0,01409 | 0,4189 |

Unmodified wild type
| Insertion into WT | Sectored | Allelic | Loss |
| --- | --- | --- | --- |
| *URA3* proxy | 0,4499 | 0,1312 | 0,3609 |

